# PLD2-phosphatidic acid recruit ESCRT-I to late endosomes for exosome biogenesis

**DOI:** 10.1101/2020.11.25.398396

**Authors:** Antonio Luis Egea-Jimenez, Stéphane Audebert, Monica Castro-Cruz, Jean-Paul Borg, Guido David, Luc Camoin, Pascale Zimmermann

**Author notes:** Correspondence and material requests should be addressed to Pascale Zimmermann or.

## Abstract

It is important to understand the biogenesis of exosomes, extracellular vesicles of endosomal origin controlling cell-to-cell communication. We previously reported that Phospholipase D2 (PLD2) supports late endosome (LE) budding and the biogenesis of syntenin-dependent exosomes. Here, we reveal that PLD2 has a broader generic effect on exosome production. Combining gain- and loss-of-function experiments, proteomics, microscopy and lipid-binding studies with reconstituted liposomes mimicking LE, we show that: (i) PLD2 activity controls the recruitment of MVB12B to LE and the exosomal secretion of ESCRT-I; (ii) loss-of-MVB12B phenocopies loss-of-PLD2, similarly affecting LE budding, the number of exosomes released and exosome loading with cargo; (iii) MVB12B MABP domain directly interacts with phosphatidic acid, the product of PLD2. We therefore propose that PLD2 and phosphatidic acid support ESCRT-I recruitment to LE for the formation of exosomes. This work highlights a major unsuspected piece of the molecular framework supporting LE and exosome biogenesis.

## Introduction

Extracellular vesicles (EVs) are lipid bilayer-limited vesicles released by any type of cell. Due to their ability to transport biologically active molecules such as cytosolic and membrane proteins, lipids and various types of nucleic acids, EVs emerge as organelles that mediate cell-to-cell communication over short and long distances. EVs playing crucial roles in homeostasis and systemic diseases, currently nearly all biomedical disciplines try to exploit EVs in diagnostics and therapeutics (Kalluri and LeBleu, 2020; Yanez-Mo et al., 2015). Yet the rational usage of EVs in biomedicine still faces many challenges (Margolis and Sadovsky, 2019) and the field awaits, among others, a better understanding of the molecular mechanisms supporting EV biogenesis.

EVs circulating n biofluids are diverse in composition, and sub-populations can be separated from each other by various methods (Thery et al., 2018). Besides apoptotic bodies, two main types of EVs are actively secreted by cells, namely microvesicles (MVs) and exosomes. MVs originate from the cell surface upon plasma membrane budding and have diameters that vary from 50 to 1,000 nm. Exosomes have the same topology as MVs but derive from endosomal membranes. They originate from the intraluminal vesicles (ILVs) that accumulate inside late endosomes or multivesicular bodies (MVBs). Upon MVB fusion with the plasma membrane ILVs are secreted in the extracellular environment, where these small vesicles (40-120 nm) are then designated as “exosomes”. Even if exosomes have been most investigated, all types of EVs have been shown to convey functional activities. Although such remains to be formally proven, exosomes might however be more potent and specific biological agents, as they are nanosized, highly stable, able to cross the blood brain barrier and contain cargo subject to double membrane sorting.

The mechanisms that govern exosome biogenesis appear diverse and might even vary between cell types (Mathieu et al., 2019; van Niel et al., 2018). The endosomal sorting complex required for transport (ESCRT) is the best-described mechanism for the formation of ILVs and MVBs (Hanson and Cashikar, 2012; Henne et al., 2011). ESCRT consists of four multimeric complexes (ESCRT-0, -I, -II and −III) and associated proteins (e.g., ATPaseVPS4, VTA1 and ALIX) assembled in an orderly manner at the endosome. ESCRT-0, I and II recognize and sequester ubiquitinated membrane proteins; ESCRT-III is implicated in membrane deformation and vesicle abscission; and the VPS4 complex is required for the recycling of ESCRT components and controls neck constriction and vesicle abscission (Christ et al., 2017). Also lipids participate in ILV formation, supporting membrane-deformation processes (Babst, 2011), as demonstrated for ceramide, the reaction product of the neutral sphingomyelinase 2 (nSMase2) (Trajkovic et al., 2008). Ceramide-induced aggregation of lipid microdomains leads to the budding of intraluminal vesicles possibly promoted by its cone-shaped structure (Marsh and van Meer, 2008). Furthermore, it is well documented that phosphoinositides participate in the recruitment of ESCRT subunits to endosomal membranes, and, thereby, regulate ILV and MVB formation in an ESCRT-dependent manner (Katzmann et al., 2003; Slagsvold et al., 2005; Teo et al., 2006; Whitley et al., 2003). Of note, such lipid-initiated recruitments and ESCRT-supported mechanisms of ILV formation were mostly studied in the context of pathways that lead to lysosomal degradation. Previous studies have also shown that phospholipase D2 (PLD2), a downstream effector of the small GTPase ARF6, is important for the production of EVs and in particular for the subclass of exosomes that involves the syntenin-pathway (Ghossoub et al., 2014; Laulagnier et al., 2004). However, the molecular mechanisms supporting PLD2 function in such events remain unknown (Egea-Jimenez and Zimmermann, 2018). PLD2 catalyzes the hydrolysis of phosphatidylcholine (PC), the most abundant membrane phospholipid, to generate phosphatidic acid (PA) and choline. Here, we identify the ESCRT-I subunit MVB12B as a prime candidate for PLD2-initiated and PA-mediated endosomal membrane targeting. MVB12B proves to be a key ESCRT-I component for ILV budding that leads to exosome formation in MCF-7 cells. We show that MVB12B accumulation in late endosomal and exosomal compartments depends on PLD2 activity and is due to a high affinity and specific interaction of the MABP domain of MVB12B with PA. We thus reveal a PLD2 actionable lipid-mediated membrane targeting of ESCRT-I important for the generation of MVBs and thereby clarify the mechanisms supporting LE and exosome biogenesis.

## Results

### PLD2 is a major stimulator of exosomal secretion, pending its phospholipase activity

First, we aimed to delineate the impact of PLD2, and PLD2 activity in particular, on microvesicles and exosomes. We therefore fractionated the conditioned medium of MCF-7 cells by differential ultracentrifugation and analyzed both the 10K fraction (enriched in microvesicles, further referred to as MVs) and the 100K fraction (enriched in exosomes, further referred to as exosomes) by particle tracking analysis (NTA). We used two independent siRNAs (siPLD2#2 and siPLD2#4), both validated for effectiveness by qRT-PCR (Fig. S1A). Both siRNAs left the number of microvesicles (MVs) unaffected (Fig. S1B left), while they decreased the number of exosomes by circa 60% (Fig. 1A left). None had an effect on exosome size (Fig. S1C left) or on cell viability (Fig. S1D). We also tested for the effect of VU0364739, a selective inhibitor of PLD2 (Lavieri et al., 2010). Clearly, the effects of lipase inhibition mimic the effects observed upon loss-of-expression, the inhibitor having no impact on the number of MVs (Fig. S1B right), the size of exosomes (Fig. S1C right) or cell viability (Fig. S1E) while decreasing the number of exosomes by also circa 60 % (Fig. 1A right). PLD2 is a multi-domain protein (Fig. 1B) containing two HKD domains supporting the catalytic activity. Overexpression of wild-type PLD2 significantly increased the number of exosomes, but two different catalytically dead forms of the protein, each with a different mutant HDK domain, did not (Fig. 1C). Cell viability was not affected after wild-type or mutant PLD2 overexpression (Fig. S1F). These data indicate that PLD2, by virtue of its lipase activity, must be major stimulator of the formation of extracellular vesicles originating from endosomes.

**Figure 1.**
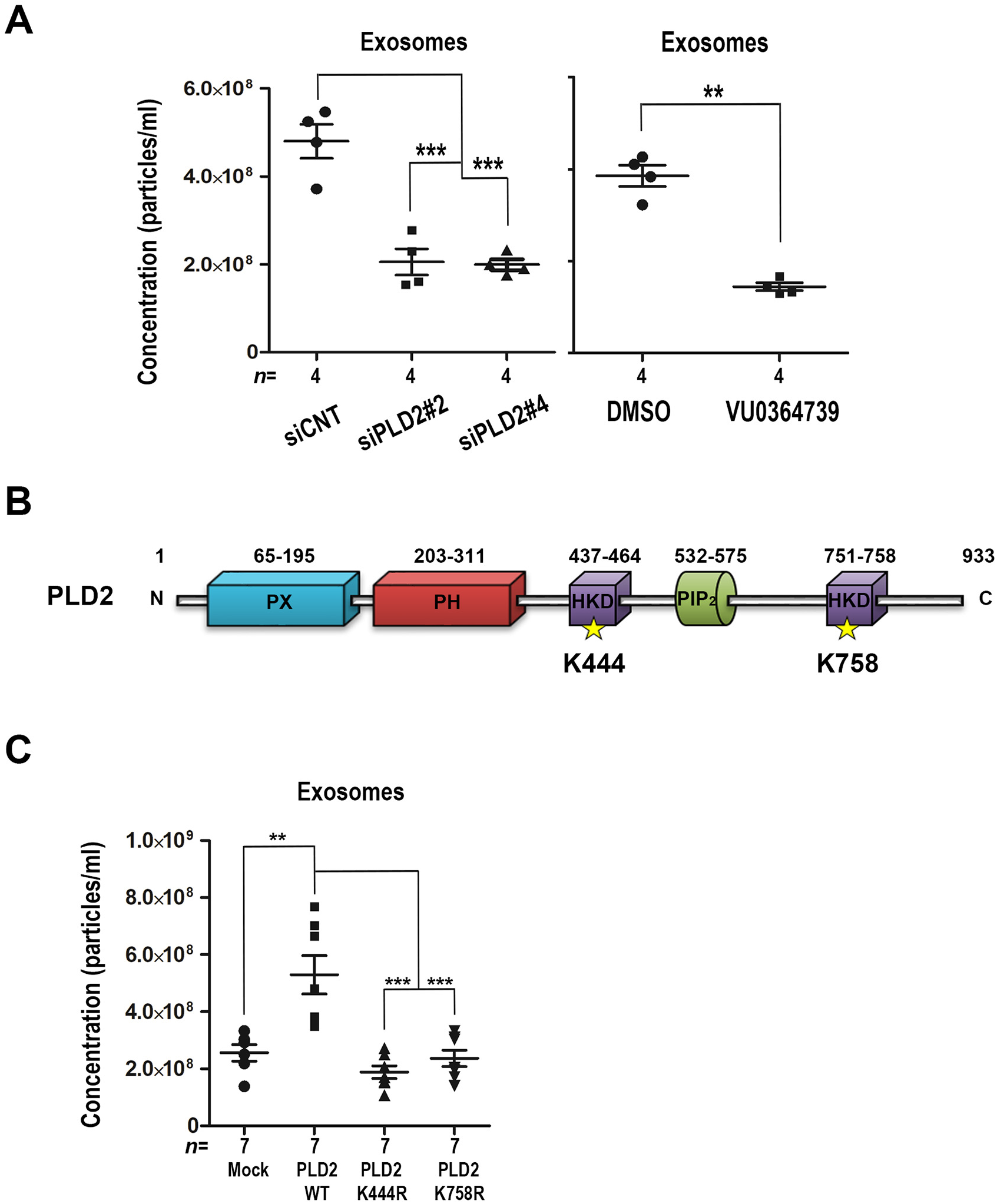
PLD2 lipase activity regulates exosome numbers. **(A)** PLD2-loss effects on the number of exosomes, studied by NTA. Exosomes, isolated by differential ultracentrifugation (100K pellet), derived from control (siCNT) and PLD2-depleted MCF-7 cells, treated with two different PLD2-specific RNAi sequences (siPLD2#2 and siPLD2#4) (left panel), or derived from controls (DMSO) and cells treated with a specific PLD2 inhibitor (VU0364739) (right panel). The dot plot represents the total number of particles, with each independent experiment represented by a different symbol (**P<0.01, ***P<0.001, Student’s t-test). Calculated for *n* independent experiments. NTA data show that PLD2-depletion and PLD2-inactivation affect the number of exosomes secreted by the cell. **(B)** Domain structure of mammalian PLD2. The point mutations of PLD2 tested in this study are represented by yellow stars. UniProt code: PLD2 014393. **(C)** PLD2-gain effects on the number of exosomes. Exosomes secreted by MCF-7 cells overexpressing *wild-type* or lipase-dead (K444R or K758R) mutant PLD2, studied by NTA. The overexpression of *wild-type* PLD2 increases the number of exosomes secreted, whereas the lipase-dead mutants do not have such effect. The dot plot represents the total number of particles, with each independent experiment represented by a different symbol (**P<0.01, ***P<0.001, Student’s t-test). Calculated for *n* independent experiments. See also Figure S1.

To complement these experiments, we investigated the effects of PLD2 loss-of-expression, inhibition of enzymatic activity, and gain-of-expression on the cellular and secreted levels of a selected set of marker proteins, commonly found in EVs and enriched in exosomes in particular. PLD2 depletion did not affect the cellular levels of ALIX, flotillin-1, syntenin, CD81, CD9 or CD63, nor did the inhibition of PLD2 activity (Fig. S2A-D). Yet, the levels of all these proteins were severely reduced in the exosomal fraction (Fig. 2A-D). Overexpression of wildtype PLD2 had also no effects on the cellular levels of these marker proteins (Fig. S2E-F), but increased their secretion in exosomes by a factor 2 to 5 (Fig. 2E-F). Overexpression of lipase-dead PLD2 yielded results similar to controls (Fig. S2E-F and Fig.2E-F), while expression of PLD2 forms mutated in residues non-conserved during evolution (Fig. S3A) gave results similar to wild-type PLD2 (Fig. S3B-D).

**Figure 2.**
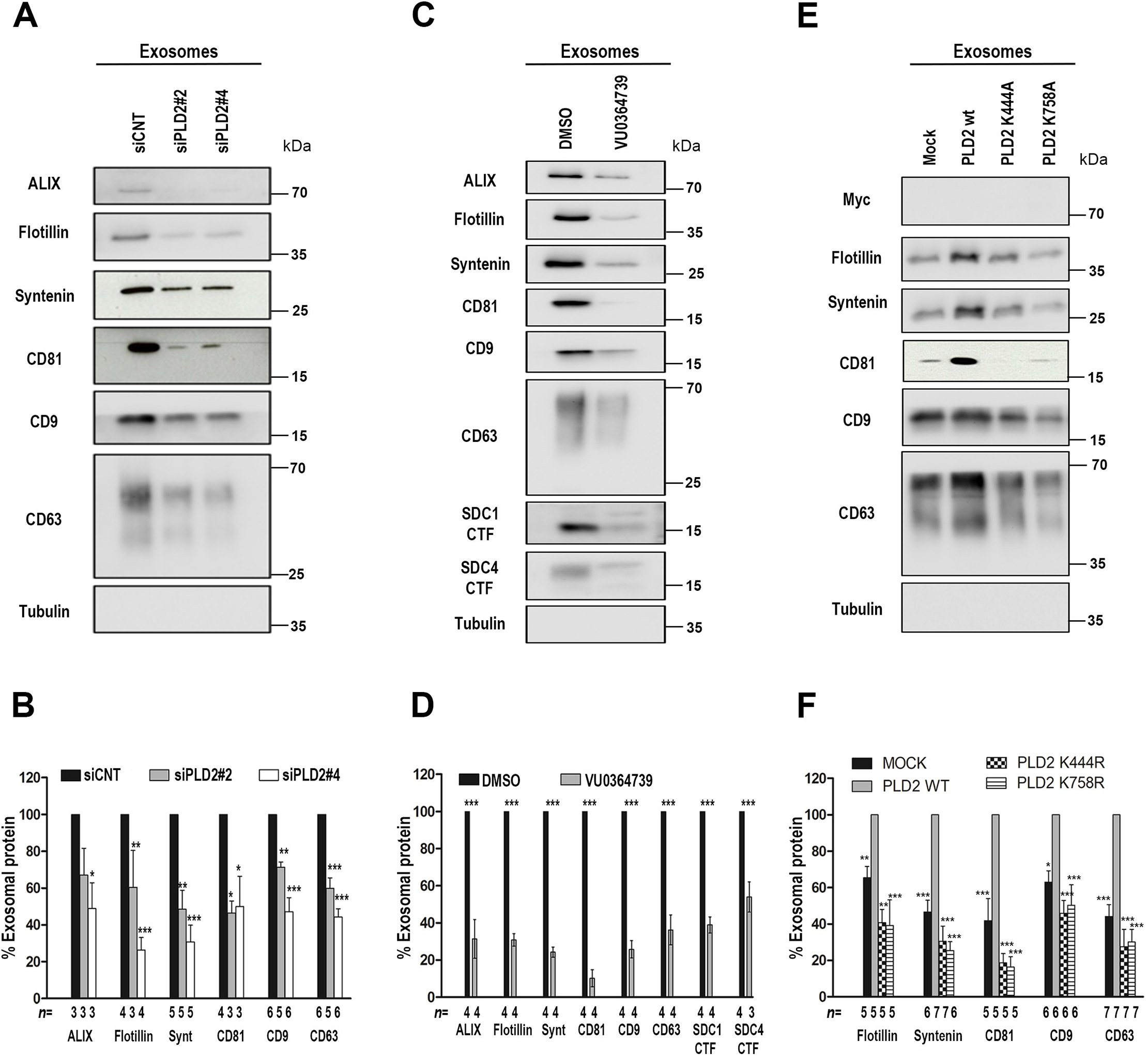
PLD2 lipase activity regulates exosome cargo. **(A-F)** PLD2 effects on the composition of exosomes, analyzed by western blotting. Antibodies for several different markers, commonly found in EVs and enriched in exosomes in particular, are used as indicated. (**A,B**) exosomes derived from control (siCNT) and PLD2-depleted MCF-7 cells, treated with two different PLD2-specific RNAi sequences (siPLD2#2 and siPLD2#4); **(C,D)** exosomes derived from controls (DMSO) and cells treated with a specific PLD2 inhibitor (VU0364739); **(E,F)** exosomes derived from MCF-7 cells overexpressing Myc-tagged *wildtype* or lipase-dead (K444R or K758R) mutant PLD2, and the respective control cells (transfected with empty vector). Histogram bars represent signal intensities, mean ± SD, relative to signals measured in control samples (black bars) (**B,D**) or in gain of *wild-type* PLD2 samples (grey bars) (**F**), taken as 100%. Statistical tests were performed using GraphPad Prism 5 software. *P<0.05, **P<0.01, ***P<0.001 (Student’s t-test). Calculated for *n* independent experiments. See also Figure S2 and S3.

MVBs can either fuse with the plasma membrane, liberating the vesicles accumulated in the endosomal lumen as exosomes, or, alternatively, with lysosomes, where their intraluminal vesicles and associated cargo are degraded. To address whether PLD2 mainly acts as an inhibitor of ‘degradative’ MVBs (with possibly more ILVs left for exosomal secretion) or a stimulator of ‘secretory’ MVBs, we investigated the effect of PLD2 inhibition under conditions where lysosomal degradation is inhibited. We reasoned that if the first scenario was taking place PLD2 inhibition would have no impact on exosomal secretion when lysosomes are inhibited, while in the second scenario PLD2 inhibition should still decrease exosomal secretion. Clearly, lysosomal inhibition did not prevent a marked decrease in the number of secreted exosomes when PLD2 is inhibited (Fig. 3A). Moreover, lysosomal inhibition did not counteract the drop of marker secretion (i.e. ‘rescue’ exosomal secretion) upon PLD2 inhibition (Fig. 3B-D). Strikingly, when PLD2 is inhibited, chloroquine still enhances the cellular levels of syntenin, CD63 and SDC4-CTFs, and this to near similar levels in cells treated with inhibitor and control cells treated with DMSO, the vehicle used for administering the inhibitor. This suggests that upon PLD2 inhibition, and thus in the near absence of exosome production, these proteins are still degraded in lysosomal compartments, in near unchanged quantities. At least for syntenin, this finding implies continued endosomal budding or membrane translocation in the absence of PLD2, to compose cargo of ILVs or autophagosomes that will be degraded in lysosomes.

**Figure 3.**
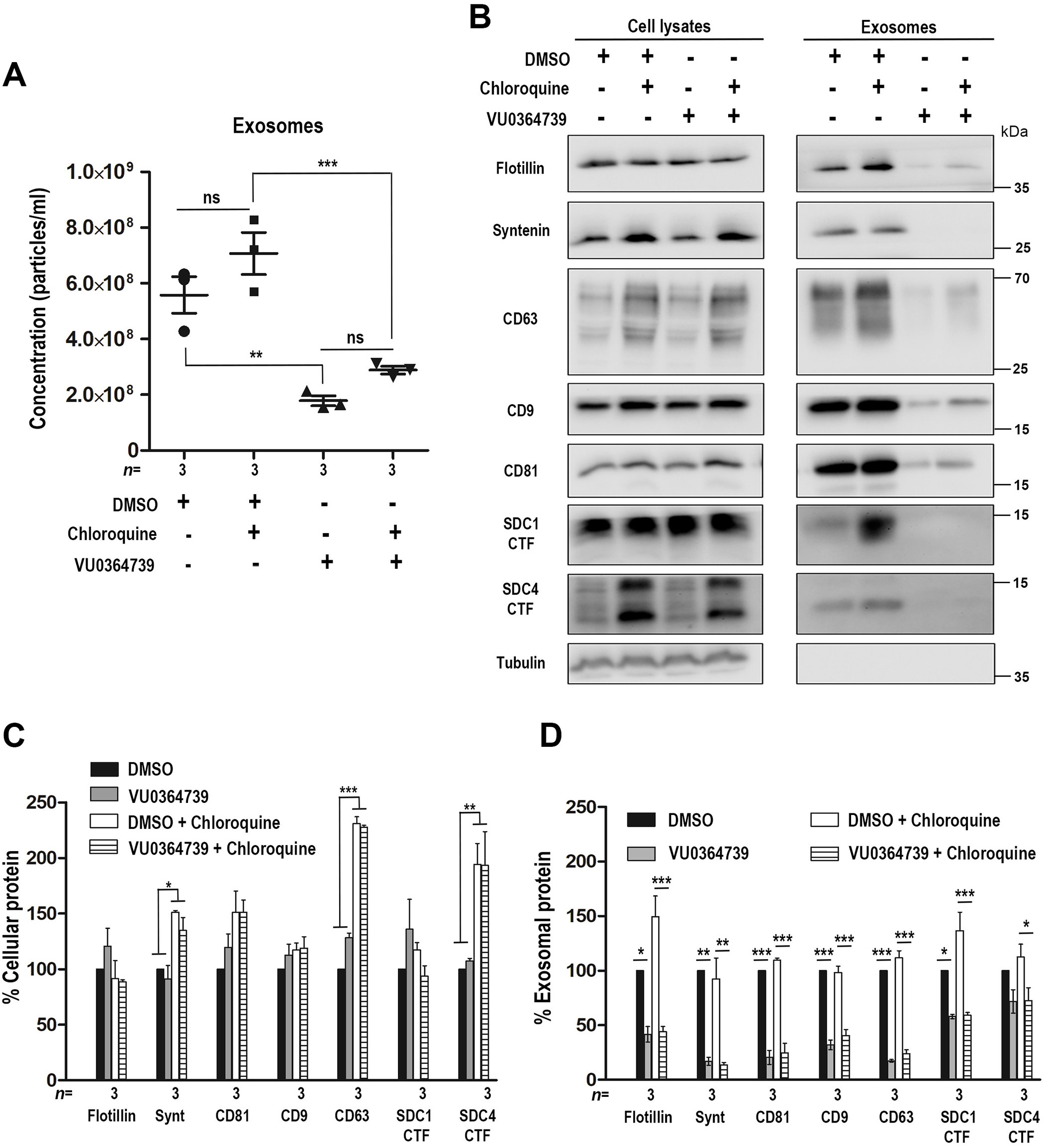
PLD2 lipase activity affects secretory MVBs and not degradative MVBs. Exosomes were purified by differential centrifugation from the conditioned media of MCF-7 cells treated with or without VU0364739 and chloroquine, as indicated, used for PLD2 and lysosome inhibition respectively. **(A)** NTA results indicate that PLD2-inactivation decreases the number of eoxomes secreted by the cell and that the combined use of chloroquine, inhibiting lysosomes, does not prevent a negative impact of PLD2 inhibition on exosome secretion. **(B)** Total cell lysates (left panel) and the corresponding exosomes (right panel) analyzed by western blotting, testing for several different markers, commonly found in EVs and enriched in exosomes in particular, as indicated. Note that while lysosome inhibition does not rescue the secretion of exosomal markers (see A), chloroquine increases the cellular levels of the markers in PLD2-inhibited cells, indicative of ongoing lysosomal degradation of these markers upon PLD2 inhibition. Histogram bars represent mean signal intensities ± SD in cell lysates **(C)** or exosomes **(D)** derived from cells treated with inhibitors, relative to signals in cell lysates or exosomes derived from DMSO-treated cells (black bars), taken as 100%. Statistical tests were performed using GraphPad Prism 5 software. **P<0.01, ***P<0.001 (Student’s t-test). Calculated for *n* independent experiments.

Taken together, these data indicate that, in MCF-7 cells, PLD2 lipase activity is a major stimulator of the secretion of exosomes and of a plethora of exosomal markers, while leaving lysosomal degradation unaffected. This suggests a generic supportive role of PLD2 in the formation and maturation of secretory MVBs and the vesicular traffic originating from these compartments.

### PLD2 supports the secretion of ESCRT-I in exosomes

In an effort to understand how PLD2 and in particular PLD2 lipase activity could so broadly impact on exosome secretion and exosomal marker release, we conducted differential proteomics experiments, comparing the exosomal fractions secreted by control cells and cells treated with siRNAs targeting PLD2 (siPLD2#2 and siPLD2#4) or PLD2 inhibitor (VU0364739). Compared to controls, 112 and 312 proteins were significantly decreased in exosomes originating from respectively siPLD2#2 and siPLD2#4 treated cells (Fig. S4A-B), while 176 proteins were decreased upon PLD2 inhibition (Fig. S4C). Among these proteins, 33 were common to the three inhibitory treatments (Fig. S5). As expected, and confirming the validity of the proteomic experiments, proteins like syntenin (SDCBP) and also CD63, CD9, ALIX (PDCD6IP) figured amongst the differentially secreted proteins (Fig. 4, in blue, in Fig. S5). More importantly, the analysis revealed that the exosomal secretion of ESCRT-I, but not that of ESCRT-0 or ESCRT-II components was significantly impaired when PLD2 function is inhibited (Fig. 4B, S4, red in S5). ESCRT-I subunits such as MVB12A and VPS37B, and the ESCRT accessory protein VPS4A were affected upon PLD2 RNAi treatment but not upon the inhibition of PLD2 activity (Fig. 4B, S4, S5B). Strikingly, TSG101, VPS28, VPS37C and MVB12B, i.e. 4 subunits participating in the assembly of ESCRT-I complexes, were present in the group of 33 proteins affected by the three inhibitory treatments (Fig. 4B, yellow). Taken together these data indicate that PLD2 and PLD2 lipase activity are important regulators of the secretion of ESCRT-I in exosomes.

**Figure 4.**
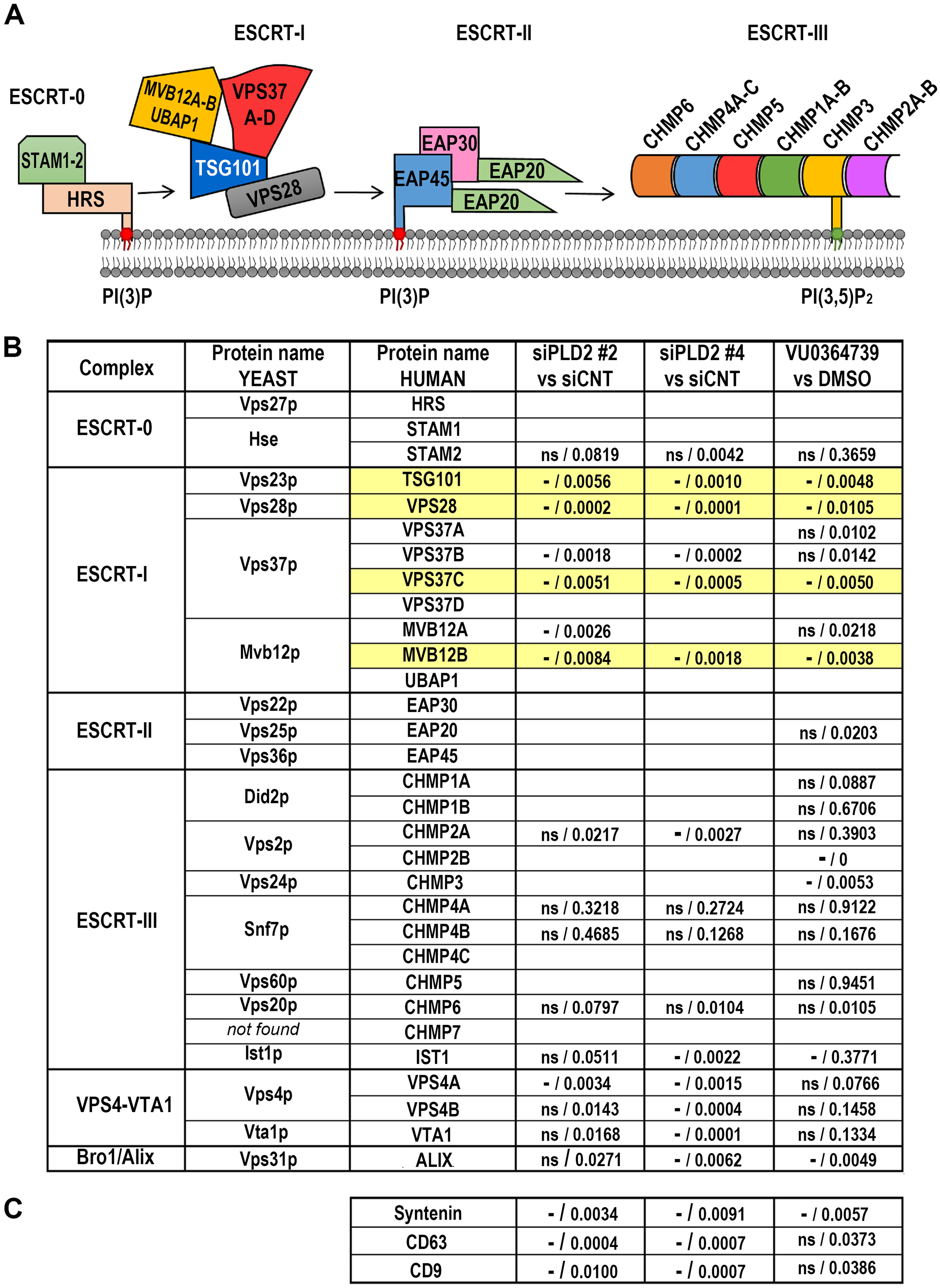
Protein subunit components of the ESCRT machinery. **(A)** Schematic representation of lipid recognition by ESCRTs on endosomal membranes. The FYVE domain of the HRS subunit of ESCRT-0 and the GLUE domain of the EAP45 subunit of ESCRT-II can interact with phosphatidylinositiol 3-phosphate (PI(3)P), a phosphoinositide enriched in early endosomal compartments, whereas the ESCRT-III subunit CHMP3 can interact with phosphatidylinositiol 3,5-bisphosphate (PI(3,5)P_2_), a phosphoinositide enriched in late endosomal compartments. Note that a role of direct lipid membrane targeting of ESCRT-I has never been described. **(B, C)** The impact of PLD2 on the composition of exosomes, as studied by proteomics. The table represents the yeast and human nomenclatures of the ESCRTs subunits, their occurrence (as detected here) in the exosomes of MCF-7 cells, and the impact of PLD2 depletion or inhibition on these levels. Exosomes originating from cells after PLD2-depletion (siPLD2 #2 and siPLD2 #4) or inactivation with a specific PLD2 inhibitor (VU0364739) were purified and analyzed by differential label-free proteomics, comparing these with their respective controls (treated with siCNT or DMSO). Proteins decreased in exosomes are represented with “—” (statistical significance p<0.01). Proteins that are detected, but are less or not significantly affected are represented with “*ns*”. Q-values are indicated in both cases. Empty boxes correspond to proteins not detected in exosomes. See also Figure S4 and S5.

### PLD2 activity controls the localization of MVB12B at late endosomes

To identify direct effectors of PLD2 activity, we performed subcellular localization studies in cells overexpressing mCherry- (Fig. 5A) or HA-tagged (Fig. S6A) ESCRT proteins. The 6 ESCRT-I proteins MVB12B, MVB12A, TSG101, VPS28, VPS37B and VPS37C, the ESCRT-III component CHMP2A and the ESCRT accessory components VPS4A and VTA were included in this study. The localizations of MVB12A, VPS28, VPS37C, VPS4A and CHMP2A were globally diffuse with both tags, even if HA-tagged VPS4A concentrates, to some extent, at the plasma membrane. VPS37B yielded a diffuse signal when tagged with mCherry, but a dotty signal when tagged with HA. TSG101 and VTA1 displayed a dotty pattern, with both tags. MVB12B concentrated at the plasma membrane and in intracellular structures irrespectively of the tag used. Noteworthy, treatment of the cells with the PLD2 inhibitor delocalized MVB12B from the intracellular structures, but not from the plasma membrane (Fig. 5A, left micrographs). The PLD2 inhibitor had no major influence on the localizations of the other ESCRT proteins (Fig. 5A). Consistent with previous data (Boura and Hurley, 2012), the intracellular structures enriched in MVB12B co-localized with the small GTPase Rab7 and thus correspond to late endosomes (Fig. 5B). PLD2 inhibition treatment did not affect Rab7 localization (Fig. S6B). These data indicate that amongst the ESCRT-I components, MVB12B is recruited to late endosomes depending on PLD2 activity.

**Figure 5.**
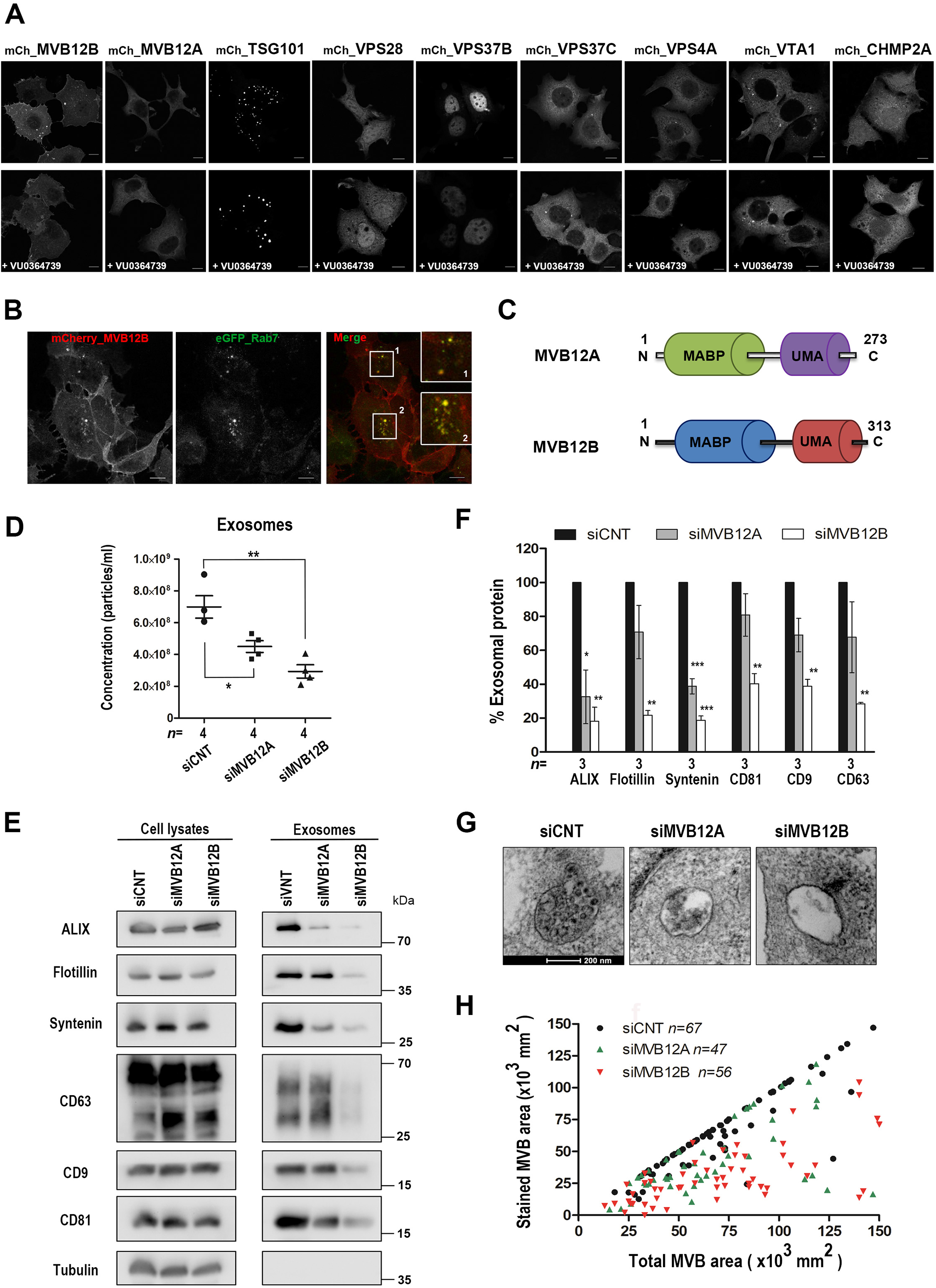
PLD2 activity directs MVB12 to late endosomes and exosomes. **(A)** Selected ESCRT proteins identified in our proteomics screen were tagged with m-Cherry (mCh_) and expressed in cells for subcellular localization studies. Representative confocal micrographs of MCF-7 cells showing the distribution of ESCRT proteins in the absence (upper line) or in the presence (lower line) of PLD2 specific inhibitor (VU0364739), as indicated. Note that the specific PLD2 inhibitory treatment de-localizes MVB12B from punctuate structures. Scale bar, 10 μm. **(B)** Intracellular MVB12B accumulates in the late endosomal compartment. Representative micrographs of MCF-7 cells transiently transfected with m-Cherry_MVB12B and eGFP_Rab7 as indicated. Note the striking co-localization of MVB12B and Rab7 in punctuate structures. Scale bar, 10 μm. **(C)** Domain structures of MVB12A and MVB12B. UniProt codes: MVB12B Q9H7P6, and MVB12A Q96EY5. **(D-F)** MVB12A and MVB12B effects on the number and composition of exosomes, studied by NTA **(D)** and western blot **(E)**, respectively. Exosomes derived from MCF-7 cells treated with a pool of four RNAi for MVB12A (siMVB12A) or MVB12B (siMVB12B) or with non-targeting control RNAi (siCNT), as indicated, were isolated by ultracentrifugation. The dot plot **(D)** represents the total number of particles, with each independent experiment represented by a different symbol. Statistical tests were performed using GraphPad Prism 5 software. *P<0.05, **P<0.01 (Student’s t-test). Calculated for *n* independent experiments. NTA results indicate that MVB12A or MVB12B depletion affects the number of Exosomes secreted by the cell, by 40 and 60% respectively. Total cell lysates (left panel) and the corresponding exosomes (right panel) were analyzed by western blotting **(E)**, testing for various markers as indicated. The histogram **(F)** represents mean signal intensities ± SD in exosomes, relative to signals measured in control sample (siCNT) (black bars), taken as 100%. Statistical tests were performed using GraphPad Prism 5 software. **P<0.01, ***P<0.001 (Student’s t-test). Calculated for *n* independent experiments. **(G)** Representative electron micrographs, illustrating the morphology of the late endosomal compartment in MCF-7 cells treated with control, MVB12A or MVB12B RNAi, as indicated. Structures were revealed by peroxidase-conjugated anti-CD63 internalized for 30 min, and staining with DAB. Scale bar, 200 nm. **(H)** Endosome filling. For each peroxidase/DAB-marked endosome, the stained sectional area of the endosome is plotted against the total sectional area of that endosome. *n* is the total number of endosomes examined in three independent experiments for control and MVB12B, and two independent experiments for MVB12A. Note that MVB12B depletion affects the number and composition of exosomes more drastically than MVB12A. See also Figure S6.

### MVB12 controls the secretion of exosomes and the formation of MVBs

Because MVB12B and MVB12A are structurally related (Fig. 5C), we tested for the effect of both proteins on exosome number (Fig. 5D) and composition (Fig. 5E-F), and on the formation of CD63-positive ILVs (Fig. 5G-H). After validation of their effectiveness, by qRT-PCRs (Fig. S6C), we measured the impact of cognate siRNAs. By NTA analysis, we found that MVB12A or MVB12B depleted cells were secreting, respectively 30 and 60% less particles compared to controls (Fig. 5D). MVB12A or MVB12B depletion had no significant impact on cell viability (Fig. S6D) and did not significantly affect the cellular levels of exosomal markers such as ALIX, flotillin-1, syntenin, CD63, CD9 or CD81 (Fig. 5E left and S6E). MVB12A-depleted cells secreted significantly less syntenin and ALIX in the exosomal fraction, while other markers remained roughly unaffected (Fig. 5E right and 5F). In contrast, MVB12B-depletion drastically reduced the exosomal secretion of all markers, including ALIX, flotillin-1, syntenin, CD63, CD9 and CD81 (Fig. 5E right and 5F). Consistently, CD63^+^-ILV formation was more drastically impaired in MVB12B-depleted cells than in MVB12A-depleted cells (Fig. 5G-H). Taken together, our data indicate an important role for mammalian MVB12B in the formation and composition of exosomes and suggest that this ESCRT-I component might work as direct effector of PLD2.

### Sensing of complexliposome-embedded phosphatidic acid by MVB12B correlates with its late-endosomal localization

The binding of MVB12B to phosphatidic acid (PA), the product of PLD2 activity, converting the membrane lipid phosphatidylcholine (PC), was measured using complex liposomes mimicking the composition of the late endosomal compartment (Skotland et al., 2017; Skotland et al., 2019). Two types of liposomes were prepared, mixing 43% cholesterol, 17% PC, 16% sphingomyelin, 9% phosphatidylethanolamine, 1% ceramide with 12% phosphatidylserine plus 2% PA, or simply 14% phosphatidylserine without PA. The latter provides equally charged liposomes, used as controls to differentiate PA-specific from nonspecific binding to negatively charged lipids. Recombinant MVB12B MABP domain, previously shown to contain a lipid-binding activity (Boura and Hurley, 2012), was utilized as analyte in SPR experiments whereby either PA-containing or control liposomes were immobilized on the sensor chip. Single-cycle kinetics unambiguously established a specific PA binding (Fig. 6A) with an apparent dissociation constant of 2.9 ± 0.6 μM, after correction for “background association” on 14% PS liposomes (Fig. S7A). To further substantiate these findings, we also tested K143 mutant forms of the MVB12B MABP domain (Fig. 6B), as this residue was previously implicated in lipid-binding (Boura and Hurley, 2012). The K143A mutant could no longer sense PA in complex liposomes (Fig. 6C). Moreover, introducing K143A or K143D mutations in full-length MVB12B abolished enrichment of the protein in late endosomal compartments (Fig. 6D) and impaired its segregation in the exosomal fraction (Fig. S7B-C). Taken together, our data indicate that PLD2 activity controls the production of exosomes, by supporting the recruitment of ESCRT-I in a MVB12B-PA dependent manner on late endosomes, thereby stimulating ILV budding (Fig. 7).

**Figure 6.**
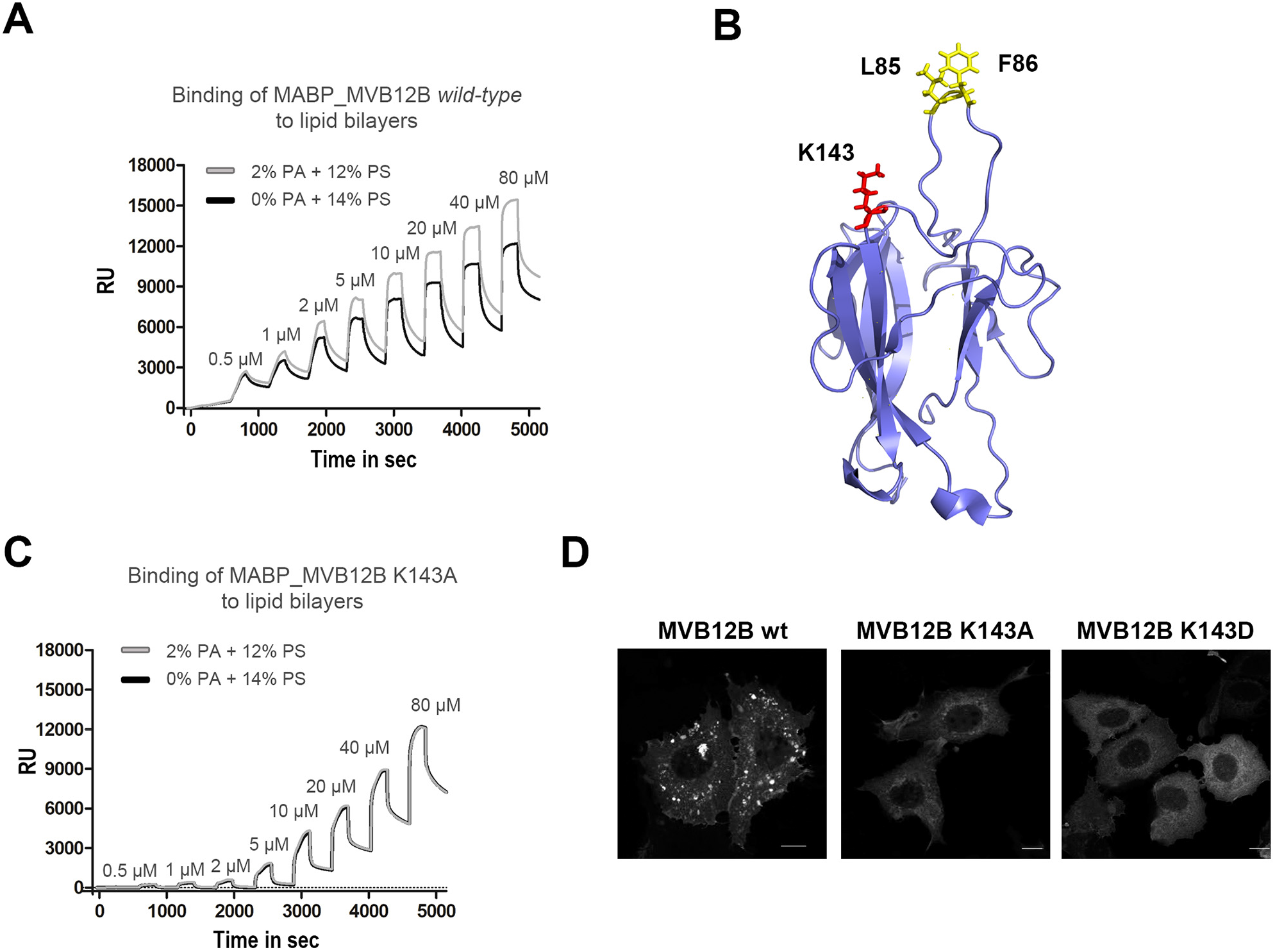
MVB12B interaction with liposome embedded PA and subcellular localization of MVB12B mutants deficient for PA binding. **(A)** Selective recognition by MABP_MVB12B of PA embedded in composite liposomes. Sensorgrams illustrating the binding of increasing concentrations of *wild-type* MABP_MVB12B to composite liposomes mimicking exosome composition, containing 43% cholesterol / 16% sphingomyelin / 17% POPC / 9% POPE / 1% ceramide and either 12% POPS / 2% PA (grey line) or 14% POPS / 0% PA (black line), the latter providing equally charged exosomes used for background reference. **(B)** Ribbon model (blue color) of the MABP domain of MVB12B. Residues implicated in lipid-binding are represented as sticks. Protein Data Bank code: MABP_MVB12B 3TOW. **(C)** Sensorgrams illustrating the binding of increasing concentrations of MABP_MVB12B K143A mutant to composite liposomes mimicking exosome compositions, as described sub A. **(D)** Confocal micrographs of MCF-7 cells overexpressing mCherry-tagged MVB12B, either *wild-type* or mutant for PA-binding, as indicated. Scale bar, 10 μm. Note the localization of MVB12B to membranes and punctate structures is impaired in PA-binding mutants. See also Figure S7.

**Figure 7.**
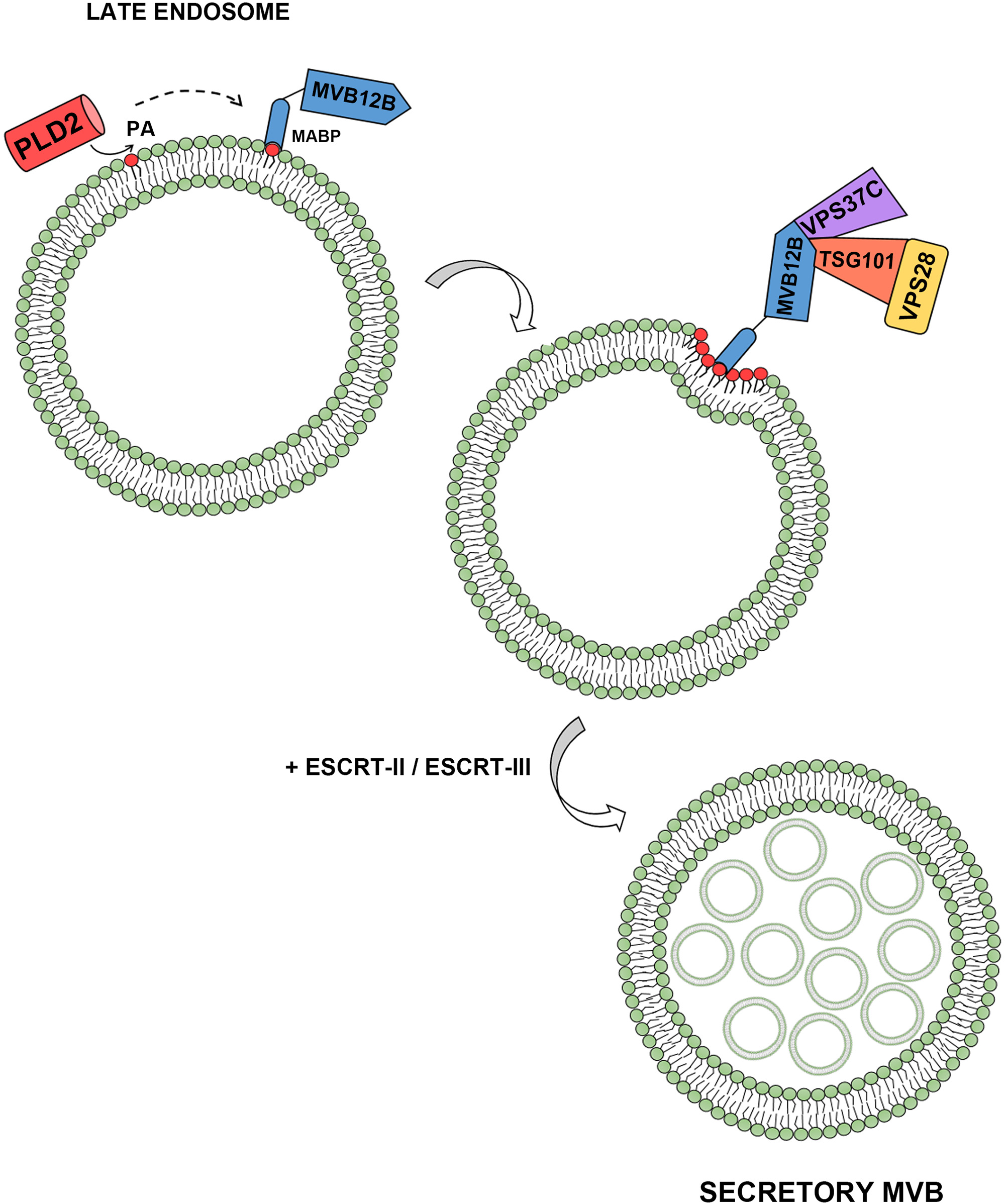
Schematic representation of ESCRT-dependent endosomal maturation triggered by PLD2 lipase activity. PLD2 activity and PA production in late endosomal compartments can support the recruitment of ESCRT-I through a direct interaction between the MABP domain of MVB12B and PA. This PLD2-dependent pathway supports the recruitment of ESCRT-I tetramers and eventually the formation of secretory MVBs.

## Discussion

In this study, we demonstrate that the catalytic activity of PLD2 is involved in endosomal budding and exosome formation. Gain-of-PLD2 boosts the production of exosomes (Fig. 1C), and increases the exosomal secretion of a wide range of marker proteins (Fig. 2E-F). Inversely, PLD2 loss-of-expression decreases the number of exosomes made (Fig. 1A) and decreases exosomal markers (Fig. 2A-D). Both PLD2-depletion and administration of the isoform-selective PLD2 inhibitor VU0364739, which - we surmise - exclusively blocks the catalytic activity without affecting other PLD2 functions, lead to similar decreases in the number and compositions of exosomes. These data confirm and extend prior data from our laboratory (Ghossoub et al., 2014) illustrating that PLD2 functions as a downstream effector of the small GTPase ARF6 in the making of syntenin-dependent exosomes. Indeed, the effects of PLD2 clearly extend beyond syntenin exosomes, as exosome cargo like CD9, CD81 and flotillin-1, previously shown to not depend on syntenin or SDC-syntenin regulators (Baietti 2012, Roucourt 2015, Imjeti 2017), are also under the control of PLD2 activity (Fig. 2, Fig. S2, Fig. S3). In contrast with its large effects on exosomes, PLD2 does not seem to participate in the generation of MVs, these remaining unaffected after either PLD2-depletion or PLD2-inactivation (Fig. S1B). Yet, it should be noted that, compared to exosomes, MCF-7 cells produce only small amounts of MVs (compare Fig. 1C with Fig. S1B). Whether PLD2 activity might control the formation of MVs in other cell types remains a possibility. Indeed, using primary alcohols to inhibit PLDs in a melanoma cell line, Muralidharan-Chari et al. observed a diminution of the shedding of MVs from the plasma membrane (Muralidharan-Chari et al., 2009). Such differences between cell types might be explained by different spatial distributions of PLD, depending on the cellular contexts. The inhibition of PLD2 blocks the secretion of exosomes even when lysosomal activity is blocked (Fig. 3A). This is indicative for a role of PLD2 in boosting secretory MVBs, rather than inhibiting degradative MVBs (Fig. 3). Of note, upon PLD2 inhibition, and thus in the near absence of exosome production, chloroquine experiments suggest that cellular proteins like syntenin, SDC4 and CD63 are still substantially degraded in lysosomal compartments (Fig. 3B-D). Also that finding is consistent with the notion that PLD2 specifically affects secretory MVBs. Yet, lysosome-inhibition does not significantly affect cellular levels of other membrane-associated exosomal cargo like CD9, CD81 and flotillin-1. This indicates that at least two (quantitatively, if not qualitatively) differential sorting mechanisms are operating for these ‘exosomal’ markers. The lysosome-limited cellular accumulations of SDC4, CD63 and syntenin might be related to their association with Tetraspanin-6 membrane webs (Ghossoub et al., 2020), while for the second, the mechanisms of clearance remain to be elucidated. Possibly CD9, CD81 and flotillin-1, composing characteristic exosome markers, nearly exclusively leave the cells via exosomal secretion, but with low turnover.

In an attempt to understand the molecular mechanisms supporting the effects of PLD2, we compared the composition of exosomes secreted by control cells, versus cells treated with (two different) siRNAs targeting PLD2 or the PLD2 inhibitor VU0364739. Label-free differential proteomics identified 33 proteins that were commonly under-represented in exosomes upon these 3 different treatments, leading us to surmise these should be the most reliable hits. Syntenin (SDCBP; q-value between 0.0034 and 0.0091, Fig. 4C) appeared in this list of ‘robust’ hits, validating the approach (Fig. S4, S5). Yet, other markers affected by a loss of PLD2, as fully validated in Western blot, like CD63, CD9 and ALIX (PDCD6IP) (Fig. 2) solely emerged in two treatments out of three (Fig. 4B, S4, S5), suggesting the criteria (q-value below 0.01) we used for the proteomics might be too stringent. In any case, amongst the 33 candidates, one striking observation was to find one complete ESCRT-I complex, namely the tetramer TSG101/VPS28/VPS37C/MVB12B as ‘missing’ (q-values between 0.0001 and 0.0101, Fig. 4B, S4). On the contrary, ESCRT-II components were not detected, while effects of PLD2 on exosomal ESCRT-0 or ESCRT-III were respectively absent or discrete (Fig. 4B, S4, S5). To our knowledge, this suggests for the first time a possible direct functional link between PLD2 and the evolutionary conserved ESCRT machinery. In yeast, ESCRT-I is represented by a single soluble hetero-tetramer consisting of VPS23/TSG101, VPS28, VPS37 and MVB12 (Kostelansky et al., 2007). In contrast to yeast, human ESCRT-I can be assembled from one out of four VPS37 isoforms (VPS37A-D) and one out of three UMA-containing proteins (MVB12A, MVB12B, and UBAP1) (Fig. 4B) (Gatta and Carlton, 2019). Thus, evolution allows in principle the formation of up to 12 different ESCRT-I complexes. Yet, it appears that the incorporation of MVB12s/UBAP1 members into ESCRT-I is highly selective with respect to their VPS37 partners (Morita et al., 2007; Stefani et al., 2011; Wunderley et al., 2014). Our proteomic analysis suggests that at least two ESCRT-I complexes might be devoted to PLD2-dependent exosome secretion, the TSG101/VPS28/VPS37C/MVB12B complex (all four subunits figuring among the robust hits), but also the TSG101/VPS28/VPS37B/MVB12A complex (VPS37B/MVB12A figuring among the 59 proteins commonly affected by siPLD2#2 and siPLD2#4) (Fig. 4B, S4, S5). In other words, our data also support the binary combination formed by VPS37C/MVB12B (Wunderley et al., 2014) and VPS37B-C/MVB12A-B (Morita et al., 2007). Interestingly, although UBAP1 mRNA levels have been identified in MCF-7 cells (Human Protein Atlas http://www.proteinatlas.org) our study does not provide evidence for the presence of VPS37A/UBAP1 (Stefani et al., 2011) in exosomes. Therefore, it might be that specific ESCRT-I complexes were devoted to exosome formation during evolution and that their assembly or recruitment is under the control of PLD2 lipase activity.

Microscopy revealed that the sub-cellular location of MVB12B is highly sensitive to PLD2 activity. Upon PLD2 inhibition the distribution of MVB12B completely changed, from a punctuate pattern into a diffuse signal (Fig. 5A). Strikingly, loss-of-function of MVB12B completely phenocopies PLD2 loss-of-function in terms of effects on ILV formation, cargo-loading and exosome number, strongly supporting a role for MVB12B as the effector of PLD2 (Fig. 5D-H). The loss-of-function of MVB12A, although structurally related to MVB12B (Fig. 5C), has milder effect on ILV formation and exosome number (Fig. 5D, G-H). Intriguingly, MVB12A appears to be cargo-selective to some extent, affecting e.g. syntenin and ALIX exosomal secretion while leaving exosomal CD9 and flotillin-1 unaffected (Fig. 5E-F). These data indicate that MVB12A and MVB12B are not functionally equivalent and that MVB12A-supported ESCRT-I complexes are most probably not strictly under PLD2 control. When overexpressed MVB12B, but not MVB12A, co-localizes with the late endosomal marker Rab7 (Fig. 5B) consistent with a previous study (Boura and Hurley, 2012). PLD2 inhibition delocalizes MVB12B from these endosomes. Of note, PLD2 inhibition did not affect the Rab7 pattern (Fig. S6B) excluding a major importance for PLD2 in the integrity/homeostasis of late endosomes. These data imply that MVB12B is a sensor of PLD2 activity, and by extension of phosphatidic acid (PA) (the product of PLD2) on late endosomes.

MVB12 was the last ESCRT-I subunit identified, and it has been described as a critical factor for sorting a subset of MVB cargoes in yeasts (Curtiss et al., 2007; Oestreich et al., 2007). Looking for structure-function explanations, we realized that unlike UBAP1 (Pashkova and Piper, 2012; Stefani et al., 2011; Wunderley et al., 2014), the other two MVB12 ESCRT-I subunits, MVB12A and MVB12B, contain a lipid targeting domain, designated as MABP (de Souza and Aravind, 2010; Pang et al., 2019) (Fig. 5C). Interestingly, Boura and Hurley showed that MVB12A and MVB12B were necessary and sufficient to recruit the complete ESCRT-I complex to liposomes made of a mixture of phosphoinositides, phosphatidylserine and cerebrosides (Boura and Hurley, 2012). Here, we demonstrate by SPR/BIAcore that the MABP domain of MVB12B is able to interact with PA, this while PA is embedded in complex composite liposomes recapitulating the lipid composition of the exosome (Fig. 6A). Binding occurs with a high affinity and selectivity and not in a charge-dependent manner, and occurs at PA levels that can be considered in the physiological range (Voelker, 1991; Welti et al., 2002). Mutation of Lysine143 to Alanine, affecting a key residue for lipid binding described previously (Boura and Hurley, 2012), was sufficient to impair MVB12B-PA interaction (Fig. 6B-C, S7A) and also impaired the Rab7-associated and PLD2-sensitive subcellular distribution of MVB12B (Fig. 6D). Revisiting the data from Boura and Hurley in light of the present study, we thus propose that the MABP domain of MVB12B functions as a PA-sensor able to translocate the protein to PA-enriched membrane domains in late endosomal compartments. Whether PLD2 recruits a preformed ESCRT-I complex via this MVB12B MABP-PA interaction or serves to support the nucleation/formation of the complex on late endosomes remains to be investigated. Yet, when mutated for PA-interaction MVB12B fails to incorporate into exosomes (Fig. S7B-C).

ESCRT-I complex from yeast binds *in vitro* to synthetic PI(3)P-containing vesicles (Kostelansky et al., 2007) through an amino-terminal basic patch present in VPS37. An equivalent of the basic patch in yeast VPS37 is not conserved in any of the four VPS37 human forms. Therefore, in mammals, the membrane-targeting determinant of ESCRT-I might be represented by the specific MABP module within MVB12 proteins. This implies that the recruitment of ESCRT-I to endosomal membranes might not solely depend on the function of the ESCRT-0, i.e. that such is not required for the ESCRT-I complex formed by the binary association between VPS37C and MVB12B. In HRS-depleted cells, membrane-associated TSG101 or VPS28 is decreased by less than 50% (Bache et al., 2003), implying that part of the ESCRT-I complex, and presumably specific tetrameric combinations of ESCRT-I subunits, might also be associated with endosomes via molecules other than ESCRT-0. Furthermore, HRS is localized mostly in early endosomes whereas TSG101 is distributed in late endosomes (Bache et al., 2003), and thus TSG101 would appear to remain associated to endosomes for longer times than HRS during endosomal maturation. Based on our findings, two possible scenarios are considered: (i) ESCRT-I requires HRS for its initial recruitment to early endosomes, and then, PA production in late endosomal compartments is able to sustain retention of ESCRT-I on endosomes during endosomal maturation, or (ii) PA production in late endosomal compartments functions as an anchor in which ESCRT-0 might not be necessary (Fig. 7). Indeed, TSG101/ESCRT-I can be recruited to various different compartments in the cell by proteins such as CEP55 and the p6 late domain of GAG during cytokinesis and HIV budding (Carlton and Martin-Serrano, 2007). In both examples, ESCRT-0 does not play a role.

Despite a large body of literature on ESCRT machinery in MVB formation, to our knowledge, lipid-signaling was not suspected to contribute to the membrane targeting of ESCRT-I in MVB formation. This contrasts with other ESCRT complexes, mechanisms of recruitment to endosomal membranes involving direct binding to PI(3)P for ESCRT-0 (HRS subunit) (Katzmann et al., 2003) and ESCRT-II (EAP45 subunit) (Slagsvold et al., 2005; Teo et al., 2006), to PI(3,5)P_2_ for ESCRT-III (CHMP3 subunit) (Whitley et al., 2003) (Fig. 4A) and to LBPA for ALIX, an accessory member of the ESCRT machinery (Matsuo et al., 2004).

We hypothesize that the pleiotropic effects of PLD2 modulations, affecting several different exosomal populations, reflect the localization of key molecules, with generic functions in membrane bending-abscission into PA-enriched compartments. PLD2 activation and subsequent PLD2 lipase activity and PA production in late endosomal compartments might recruit the specific ESCRT-I tetramer TSG101/VPS28/VPS37C/MVB12B via the MVB12B subunit. Interestingly, it has been reported that ALIX, an auxiliary component of the ESCRT machinery, can also bind directly to the N-terminal UEV domain of TSG101 (Strack et al., 2003) which provides a direct link between ESCRT-I and ESCRT-III complexes. ESCRT-III is the complex most directly involved in reshaping and severing membranes (Schoneberg et al., 2017). In this proposed mechanism, the assembly/formation of MVB12B-positive ESCRT-I hetero-tetramers might possibly be due to a direct MVB12B-PA interaction after PLD2 activation in late endosomal compartments, whose membrane-recruitment might trigger the formation of exclusively “secretory” MVBs.

Finally, it is already known that lipids such as ceramide (Trajkovic et al., 2008) or lysobisphosphatidic acid (LBPA) (Matsuo et al., 2004) participate in exosome biology, but the exact mechanisms remain unclear. Despite its simple structure and relatively low abundance (representing 1-4% of the total phospholipids (Voelker, 1991; Welti et al., 2002), PA is an important phospholipid for membrane dynamics, i.e. controlling fission and fusion events (Kooijman et al., 2003). The function of PA is likely in part due to its ability to induce a negative membrane curvature. Because of the small headgroup of this lipid, membrane domains enriched in PA will tend to bend and ultimately form vesicles. Here we reveal that the production of PA in late endosomal compartments, supported by PLD2 activity, allows the recruitment of ESCRT-I through a MVB12B-PA direct interaction thereby recruiting ESCRT-I tetramers and producing secretory MVBs. This work highlights a major unsuspected piece of the molecular framework supporting late endosome and exosome biogenesis.

## REAGENTS AND TOOL TABLE

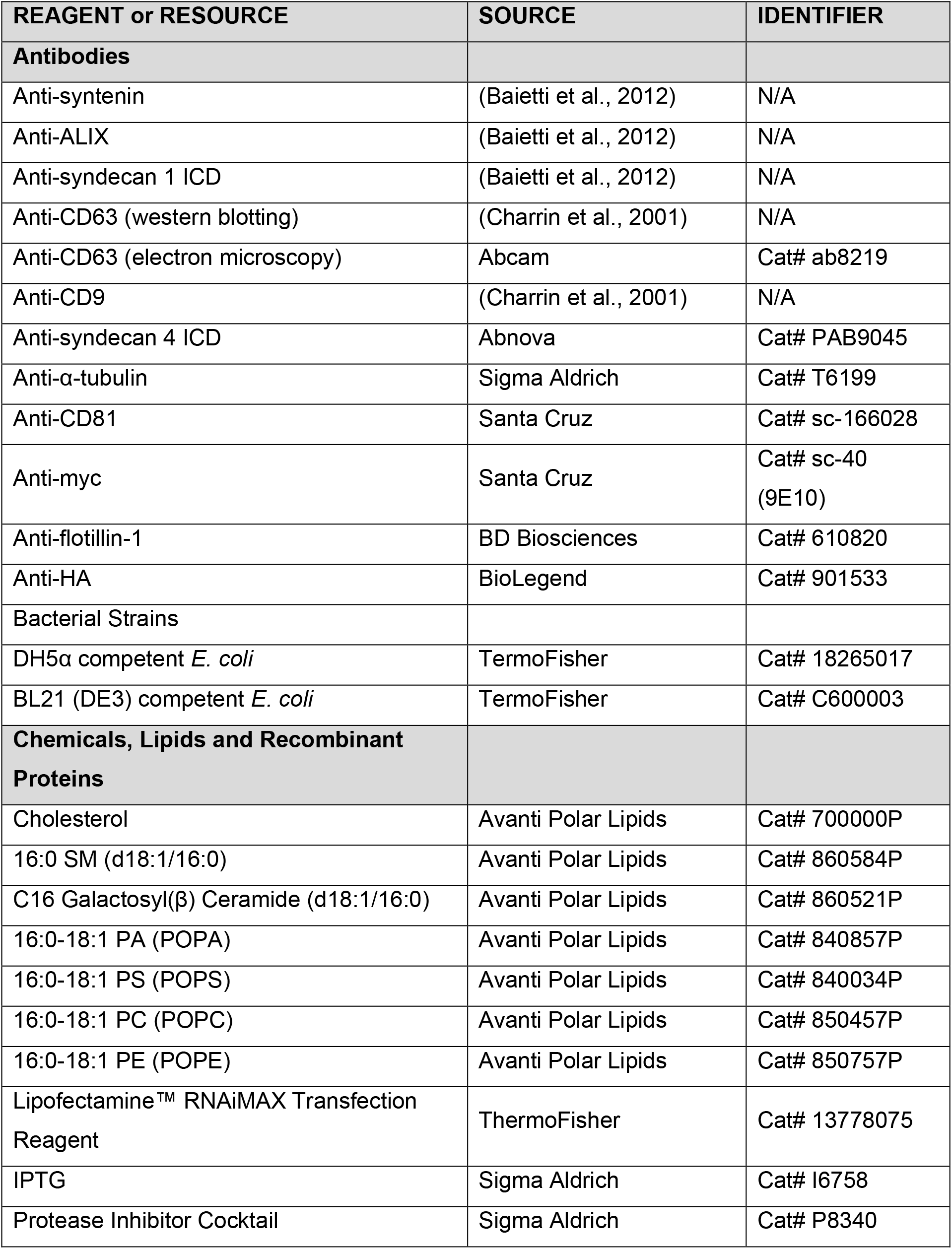
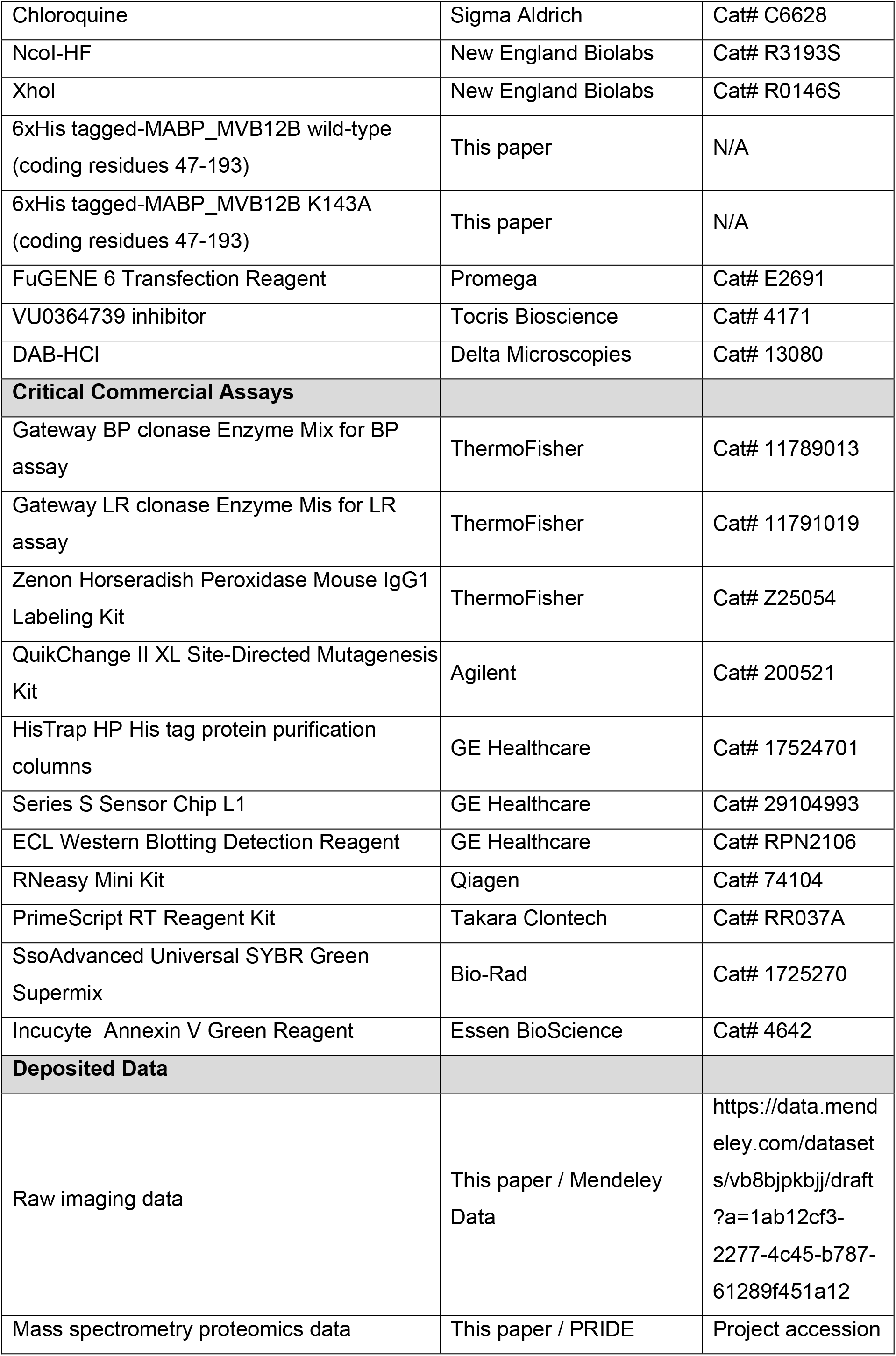
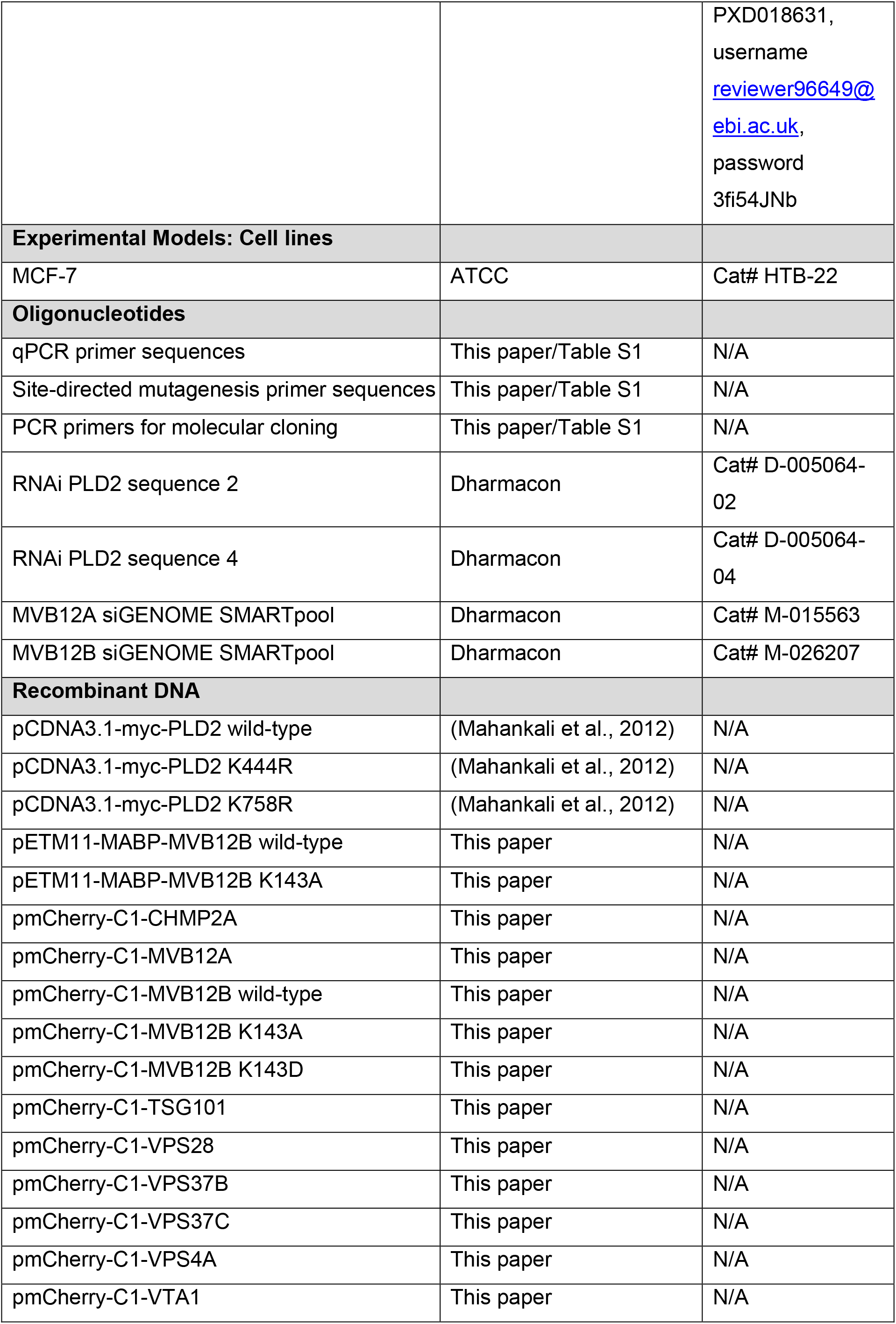
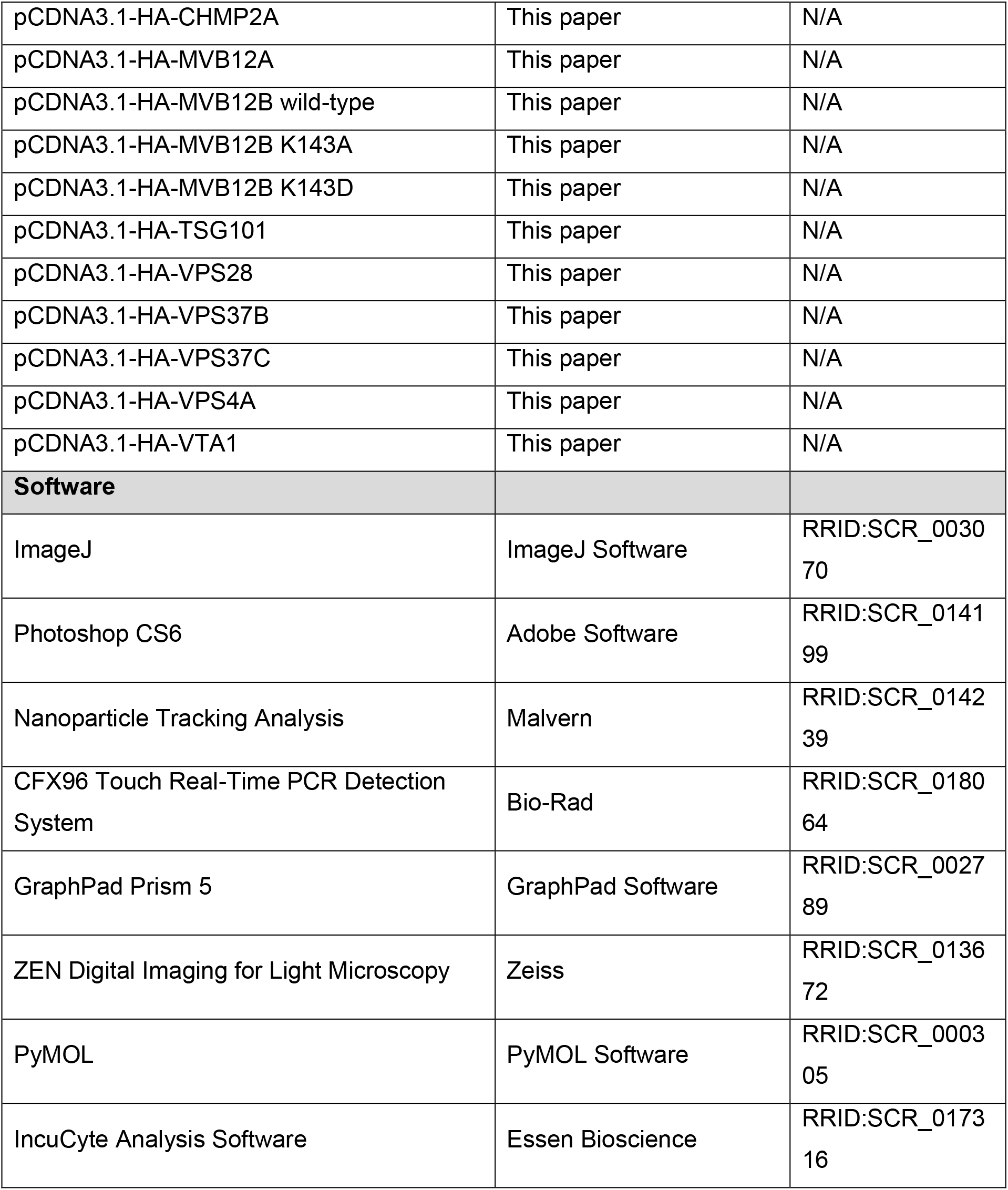

### Contact for reagents and resource sharing

Further information and requests for resources and reagents should be directed to and will be fulfilled by the lead contact, Pascale Zimmermann

### Experimental model

The MCF-7 cell line was obtained from the American Type Culture Collection (ATCC) and was routinely grown in DMEM/F12 medium (Life Technologies) supplemented with 10% fetal bovine serum (Gibco). Cell line was cultured at 37° C in a humidified incubator containing 5% CO_2_.

## Method details

### Expression vectors

Human cDNA encoding TSG101 and VPS4A was PCR amplified from a MCF-7 cDNA library using appropriate primers (Table S1) containing the two recombinant Gateway sites *attB1.1* and *attbB.1* and subcloned into a pDONR/Zeo Gateway Donor vector (Invitrogen) as the entry clones. ORF clones encoding VPS37B, VPS37C, MVB12A, MVB12B, VTA1 or CHMP2A subcloned in the pDONR223 Gateway Donor vector (Invitrogen) were obtained from Montpellier Genomic Collections (Montpellier, France). Gateway recombination reactions were performed with LR Clonase II enzyme mixes (Invitrogen) for the cloning into pmCherry-C1 and pcDNA3.1-HA vectors, with mCherry or HA respectively at the N-terminus. For the construction of the 6xHis-tagged MABP domain of MVB12B, the cDNA fragment for MABP was amplified by PCR, digested with NcoI/XhoI, and inserted into the pETM11 vector (EMBL Heidelberg). Site-directed mutagenesis was performed using QuickChange protocol (Agilent) with appropriate primers (Table S1). pCDNA3.1-myc-PLD2 *wild-type* and lipase-dead constructs were kindly provided by Dr. Julian Gomez-Cambronero, from Wright State University School of Medicine (Dayton, OH) (Mahankali et al., 2012). All the sequences of the coding region were confirmed by DNA sequencing.

### Cells experiments and reagents

Serum used for exosome purification was depleted of EVs, by prior overnight centrifugation at 140,000 x *g*. Cells were regularly controlled for the absence of mycoplasma contamination. For RNAi experiments, cells were transfected with 20 nM RNAi using Lipofectamine RNAiMAX transfection reagent (ThermoFisher) and analyzed after 72 h after RNAi treatment. All RNAi targeting human sequences (siGENOME) and non-targeting control RNAi (siCNT) were purchased from Dharmacon Inc. PLD2#2 (5-GGACCGGCCUUUCGAAGAU-3’); PLD2#4 (5’-CAGCAUGGCGGGACUAUAU-3’). MVB12A siGENOME SMARTpool (M-015563); MVB12B siGENOME SMARTpool (M-026207). For transient expressions, the cells were transfected 16 h after plating using FuGENE^®^ 6 Transfection Reagent (Promega). For PLD2 inhibition, cells were treated with 50 nM VU0364739 inhibitor (N-[2-[1-(3-Fluorophenyl)-4-oxo-1,3,8-triazaspiro[4.5]dec-8-yl]ethyl]-2-naphthalenecarboxamide) (Lavieri et al., 2010) for 16 h (Tocris Bioscience). For lysosomal inhibition, cells were treated with 100 μM chloroquine (Sigma-Aldrich) for 16 h.

### Immunofluorescence staining and confocal microscopy

For microscopic analysis, MCF-7 cells were plated on glass coverslips in a 24-well plate, fixed with 4% paraformaldehyde for 20 min, washed in PBS and then incubated with anti-myc antibody (sc-40 AC, Santa Cruz, 1:1000 dilution), anti-HA antibody (901533, clone 16B12, BioLegend, 1:1000 dilution) and appropriated Alexa-conjugated secondary antibodies (Molecular Probes) diluted in PBS containing 1% BSA and 0.5% Tween 20. Coverslips were mounted on Mowiol-DABCO and examined in a Zeiss Meta confocal microscope (LSM 510 META, Zeiss, France) with a UV laser and x 63 objective. Confocal images were analyzed using ImageJ (National Institutes of Health, Bethesda, MD) and Photoshop (Adobe, San Jose, CA) software.

### Purification of extracellular vesicles

Preliminary note: In this study, we used the term exosomes for small extracellular vesicles pelleting at 100,000 x g and microvesicles for those pelleting at 10,000 x g. This is an approximation and we recommend readers that might be confused by methodology to read the MISEV guidelines 2014 and 2018 (Lotvall et al., 2014; Thery et al., 2018). MCF-7 cells were seeded into 6-cm dishes and incubated in a 37 °C incubator containing 5% CO_2_. For comparative analyses, in gain- and loss-of-function studies, exosomes were collected from equivalent amounts of culture medium, conditioned by equivalent amounts of cells. After the required time of cDNA or RNAi treatments, the cell layers were washed twice with PBS and refreshed with DMEM/F12 containing 10% vesicle-depleted FCS. Cell-conditioned media were collected 16 h later. Exosomes were isolated from these media by three sequential centrifugation steps at 4 °C: 10 min at 500 × *g*, to remove cells; 30min at 10,000 × *g*, to remove microvesicles; and 3h at 140,000 × *g*, to pellet exosomes, followed by one wash (suspension in PBS/centrifugation at 140,000 × g), to remove soluble serum and secreted cellular proteins. Exosome pellets prepared by differential high-speed centrifugation were then re-suspended in PBS. The corresponding cell layers were washed in cold PBS and scraped on ice. Lysates from corresponding cultures were cleared by centrifugation at 300 × *g* for 5 min at 4 °C and then re-suspended in lysis buffer (30 mM Tris-HCl pH 7.4, 150 mM NaCl, 1% NP40, 10 mM EDTA and protease inhibitor cocktail dilution 1:1000 dilution, reference P8340-5ML from Sigma-Aldrich). Although little variations were observed from sample to sample (max 10%), amounts of exosomes loaded for the western blot were normalized according to the number of parent cells from where exosomes were derived.

### Preparation of liposomes

Lipid vesicles were generated by mixing chloroform solutions of cholesterol (700000P); N-palmitoyl-D-erythro-sphingosylphosphorylcholine (SM-860584P); D-galactosyl-ß-1,1’ N-palmitoyl-D-erythro-sphingosine (Cer-860521P); 1-palmitoyl-2-oleoyl-sn-glycerol (DG-800815C); 1-palmitoyl-2-oleoyl-sn-glycero-3-phosphate (PA-840857P); 1-palmitoyl-2-oleoyl-sn-glycero-3-phospho-L-serine (PS-840034P); 1-palmitoyl-2-oleoyl-glycero-3-phosphocholine (PC-850457P) and 1-palmitoyl-2-oleoyl-sn-glycero-3-phosphoethanolamine (PS-850757P) from Avanti Polar Lipids (Alabaster, Alabama, USA) in the desired proportions. Lipids were dried under a stream of nitrogen followed by exposure to high vacuum for 30 min. Dried phospholipids were resuspended in the corresponding buffer (25 mM HEPES, pH 7.4 and 150 mM NaCl) by vigorous vortexing and also using a bath sonicator to get all of the lipid into suspension. Then, the lipid suspension was subjected to eight rounds of freezing (liquid N2)-thawing (in the bath sonicator at 40 °C). The large unilamellar phospholipid vesicles of ~100nm diameter were prepared by extruding rehydrated phospholipid suspensions through two stacked 0.1 μm polycarbonate membranes (Avanti Polar Lipids).

### Protein production and purification

Competent E. coli (BL21 DE strain) cells were transformed with the human MABP domain (*wild-type* and K143A mutant) of MVB12B (coding residues 47-193) in pETM-11 vector (EMBL Heidelberg) for the expression of N-terminally 6xHis-tagged protein and then plated on LB agar plates with 50 μg/ml kanamycin at 37°C for 16 h. The culture and induction were in TYB medium (5-6 hours induction at 30°C, after addition of 0.5 mM IPTG). Fusion proteins were recovered from the cell lysates by conventional affinity chromatography on His-trapTMHP columns (GE Healthcare, 17-5247-01) using an Akta Explorer system (Amersham Pharm BiotechGE Healthcare).

### Surface Plasmon resonance experiments

The SPR experiments were at 25 °C, in a BIAcore T200 instrument (GE Healthcare). The L1 chip (GE Healthcare) was coated with composite liposomes (43% cholesterol / 16% sphingomyelin / 17% POPC / 12% POPS / 9% POPE / 1% ceramide and 2% POPA or 14% POPS / 0% POPA for background reference and equally charged liposomes). Liposomes were injected at a flow rate of 5 μL per min until 5,000 RU of immobilization was reached. A short pulse of 10 mM NaOH at a flow rate of 100 μl per min was performed to remove loose liposomes before measurements. Analytes (His-tag fusion proteins) were perfused in running buffer (25mM HEPES, pH 7.4 and 150mM NaCl) without regeneration steps between injections following the single-cycle kinetics method as was described in previous work (Egea-Jimenez et al., 2016).

### Western blotting

The proteins were heat-denaturated in Laemmli sample buffer, fractionated in 15% gels by SDS-PAGE, and electro-transferred to nitrocellulose membrane. The membranes were blocked with 3% BSA and incubated with primary antibodies. Homemade anti-syntenin, anti-ALIX and anti-syndecan-1 intracellular domain antibodies were as described earlier (Baietti et al. 2012). Anti-CD63 and anti-CD9 antibodies were provided by E. Rubinstein (Charrin et al., 2001) (Université Paris-Sud UMRS_935, Villejuif). Other antibodies were from commercial sources and used as recommended by the manufacturers: anti-syndecan-4 intracellular domain (PAB9045, Abnova, 1:2000 dilution), anti-α-tubulin (T6199, Sigma-Aldrich, 1:10.000 dilution), anti-CD81 (D-4, sc-166028, Santa Cruz, 1:1000 dilution), anti-flotillin-1 (610820, BD Biosciences, 1:1000 dilution). After washing, the membranes were incubated with HRP-conjugated secondary antibody for 1 h. Signals were visualized with enhanced chemiluminescence detection reagent (Amersham Pharmacia Biotech) and were quantified by densitometric scanning, using ImageJ (National Institutes of Health, Bethesda, MD).

### Particle size analysis using Nanoparticle Tracking Analysis (NTA)

Microvesicles and exosomes collected by ultracentrifugation were resuspended in 0.22 μM filtered PBS and analyzed at similar dilutions with a Nanosight NS-300 instrument (Malvern Instruments, UK). 10% of the secretome of approximately 5 x 10^6^ MCF-7 cells were injected for NTA analysis. All measurements were made in triplicate with flow rate of 50 μL per min applied with an automated syringe pump and at a controlled temperature of 25 °C. Three videos of 60 s each were recorded and used to calculate mean values of particle concentration and size. All analyses were performed using NTA software v3.1 (Malvern, UK).

### Electron Microscopy

For the MVB morphology studies, cells were incubated for 30 min at 37 C with an anti-CD63 antibody (Abcam) labelled with horseradish peroxidase (Z 25054 Molecular Probes). The cells were then washed with PBS and fixed for 30min in 3% glutaraldehyde in 0.1M sodium cacodylate (NaCaC) buffer. After several washes in 0.1M NaCaC, the cells were incubated with diaminobenzidine (DAB; Delta Microscopies) in 0.05M Tris-HCl, pH 7.6. The enzymatic reaction was initiated by adding H_2_O_2_ at a final concentration of 0.01% and allowed to proceed for 45 min. After washing in Tris-HCl buffer, the cells were post-fixed for 1 h with 1% osmium tetroxide in 0.1M phosphate buffer, washed, scraped, re-suspended in 2% agar and pelleted. Sliced agar blocks were dehydrated in a series of ethanol and embedded in Epon resin polymerized for 48 h at 60 C. Eighty nanometer sections (Ultracut UC7 Leica) were counterstained with 2% uranyl acetate and lead citrate (Reynold’s), and were examined with a FEI G2 (ThermoFisher, USA) at 200 kV and images were taken with a Veleta camera. MVBs sectional areas were measured using ImageJ software, and the data were analysed using Excel software.

### Cell viability assay

Live cell analysis was performed using an IncuCyte ZOOM systems (Essen BioScience) to determine the cytotoxic effects of siRNA treatment, vector expression, or PLD2 inhibition. This system allows an automated analysis method of monitoring live cells in standard incubator conditions. MCF-7 cells were seeded at confluency 5 x10^3^ in 96-well plates (Corning) and incubated with the correspondent siRNA treatments, expression vectors, or solely with complete medium. After overnight incubation, the culture medium of the siRNA and vector treated cells was replaced with 100 μL of fresh complete medium containing 1:200 green 490 - 515nm Annexin V reagent (Essen BioScience) as recommended by the manufacturer. To the MCF-7 cells initially incubated solely with complete medium, 100 μL of fresh complete medium containing the PLD2 inhibitor and 1:200 Annexin V were added. Two images were acquired per well, from three technical replicates every 2-4 h for 48 to 70 h, using a 10 X objective lens and data were analyzed using the IncuCyte Basic software.

### Mass spectrometry analysis and proteins quantification

30% of exosomes preparations were loaded on NuPAGE 4–12% bis-Tris acrylamide gels (Life Technologies) to stack proteome in a single band that was stained with Imperial Blue (Thermo Fisher Scientific) and cut from the gel. Gels pieces were submitted to an in-gel trypsin digestion (Shevchenko et al., 1996). Peptides were extracted from the gel and dried under vacuum. Samples were reconstituted with 0.1% trifluoroacetic acid in 4% acetonitrile and analyzed by liquid chromatography (LC)-tandem MS (MS/MS) using a Q Exactive Plus Hybrid Quadrupole-Orbitrap online with a nanoLC Ultimate 3000 chromatography system (Thermo Fisher Scientific™, San Jose, CA). For each biological sample, 3 microliters corresponding to 15% of digested sample were injected in triplicate on the system. After preconcentration and washing of the sample on a Acclaim PepMap 100 column (C18, 2 cm × 100 μm i.d. 100 A pore size, 5 μm particle size), peptides were separated on a LC EASY-Spray column (C18, 50 cm × 75 μm i.d., 100 A, 2 μm, 100A particle size) at a flow rate of 300 nL/min with a two steps linear gradient (2-22% acetonitrile/H20; 0.1 % formic acid for 220 min and 22-32% acetonitrile/H20; 0.1 % formic acid for 20 min). For peptides ionization in the EASYSpray source, spray voltage was set at 1.9 kV and the capillary temperature at 250 °C. All samples were measured in a data dependent acquisition mode. Each run was preceded by a blank MS run in order to monitor system background. The peptide masses were measured in a survey full scan (scan range 375-1500 m/z, with 70 K FWHM resolution at m/z=400, target AGC value of 3.00×10^6^ and maximum injection time of 100 ms). Following the high-resolution full scan in the Orbitrap, the 10 most intense data-dependent precursor ions were successively fragmented in HCD cell and measured in Orbitrap (normalized collision energy of 25 %, activation time of 10 ms, target AGC value of 1.00×10^5^, intensity threshold 1.00×10^4^ maximum injection time 100 ms, isolation window 2 m/z, 17.5 K FWHM resolution, scan range 200 to 2000 m/z). Dynamic exclusion was implemented with a repeat count of 1 and exclusion duration of 20 s.

### Mass spectrometry data processing

Relative intensity-based label-free quantification (LFQ) was processed using the MaxLFQ algorithm from the freely available MaxQuant computational proteomics platform, version 1.6.3.4. Spectra were searched against the Human database, extracted from UniProt on the 9^th^ of December 2019 and containing 20379 entries (reviewed) supplemented with 229 major bovine contaminants proteins identified in calf serum used for cells culture. The false discovery rate (FDR) at the peptide and protein levels were set to 1% and determined by searching a reverse database. For protein grouping, all proteins that cannot be distinguished based on their identified peptides were assembled into a single entry according to the MaxQuant rules. The statistical analysis was done with Perseus program (version 1.6.2.1) from the MaxQuant environment (www.maxquant.org). Quantifiable proteins were defined as those detected in above 70% of samples in one condition or more. Protein LFQ normalized intensities were base 2 logarithmized to obtain a normal distribution. Missing values were replaced using data imputation by randomly selecting from a normal distribution centred on the lower edge of the intensity values that simulates signals of low abundant proteins using default parameters (a downshift of 1.8 standard deviation and a width of 0.3 of the original distribution). To determine whether a given detected protein was specifically differential, a two-sample t-test were done using permutation-based FDR-controlled at 0.01 and employing 250 permutations. The p value was adjusted using a scaling factor s0 with a value of 0.4 for the loss-of-PLD2 (siPLD2#2 and siPLD2#4 conditions) with n=2, and a value of 2 for the PLD2 inhibition treatment (VU0364739 condition) with n=1.

### RNA extraction, reverse transcription and real-time qPCR

Total RNA was extracted from cells using Nucleospin RNA isolation kit (Qiagen, USA). 1 μg of total RNA from each condition was converted into cDNA using PrimeScript RT reagent with gDNA Eraser kit (Takara Clontech) and MJ Research PTC-220 Dyad PCR System (Conquer Scientific). Quantitative PCR reactions were performed using SsoAdvanced Universal Sybr Green Supermix kit (Bio-Rad), and analyzed using CFX96 Touch Real-Time PCR Detection System (Bio-Rad). The expression levels were normalized to the internal control actin and GAPDH gene expressions. The PCR programme included 35 cycles of denaturation at 95 °C for 15 s and 30 s of annealing and extension at 60 °C. The sequence information of qPCR primers used for the expression analysis is listed in Table S1.

## Quantification and statistical analysis

Each experiment was repeated three times or more, unless otherwise is noted. Statistical analysis was performed using the standard two-tailed Student t-test, and * P < 0.05, ** P < 0.01, *** P < 0.001 were considered statistically significant. Statistical test was performed using GraphPad Prism 5 software.

## Date and software availability

Sequence information about all the oligos used in this study can be found in Table S1. Original imaging data have been deposited to Mendeley Data at: https://data.mendeley.com/datasets/vb8bjpkbjj/draft?a=1ab12cf3-2277-4c45-b787-61289f451a12. The mass spectrometry proteomics data, including search results, have been deposited to the ProteomeXchange Consortium (http://www.proteomexchange.org/) via the PRIDE (Vizcaino et al., 2014) partner repository with the dataset identifier: Project accession PXD018631, username, password 3fi54JNb.

## Acknowledgments

We thank the PiCSL-FBI core electron microscopy facility (Aïcha Aouane, Fabrice Richard and Nicolas Brouilly, Aix Marseille Univ, CNRS, IBDM, Marseille, France), member of the national infrastructure France-BioImaging supported by the French National Research Agency (ANR-10-INBS-0004). We are grateful to Eric Bailly for the molecular biology advice. This work was supported by grants from the Agence Nationale de la Recherche (ANR-18-CE13-0017, Project SynTEV), and the Fund for Scientific Research-Flanders (FWO Grant G0C5718N). Antonio Luis EGEA-JIMENEZ was recipient of a postdoctoral fellowship of La Ligue contre le Cancer (France). This project has received funding from the European Union’s Horizon 2020 research and innovation program under the Marie Sklodowska-Curie grant agreement No 747025. Proteomic analyses were done using the mass spectrometry facility of Marseille Proteomics (marseille-proteomique.univ-amu.fr) supported by IBISA (Infrastructures Biologie Santé et Agronomie), Plateforme Technologique Aix-Marseille, the Cancéropôle PACA, Région Sud-Alpes-Côte d’Azur, the Institut Paoli-Calmettes, the Centre de Recherche en Cancérologie de Marseille, Fonds Européen de Developpement Régional and Plan Cancer. We thank Emilie Baudelet for technical assistance in mass spectrometry analysis. J-P.Borg lab is also funded by La Ligue Nationale Contre le Cancer (Label Ligue J.P.B.). J-P.B. is a scholar of Institut Universitaire de France.

## Author contributions

A.L.E-J designed experiments; A.L.E-J, S.A., M.C-C and L.C. performed experiments; J.-P.B. contributed new reagents/analytic tools; P.Z. conceived the project and supervised the research; A.L.E-J., S.A., L.C., G.D., and P.Z. analyzed data; and A.L.E-J., G.D., and P.Z. wrote the paper. All authors revised the manuscript.

---

**Figure S1.**
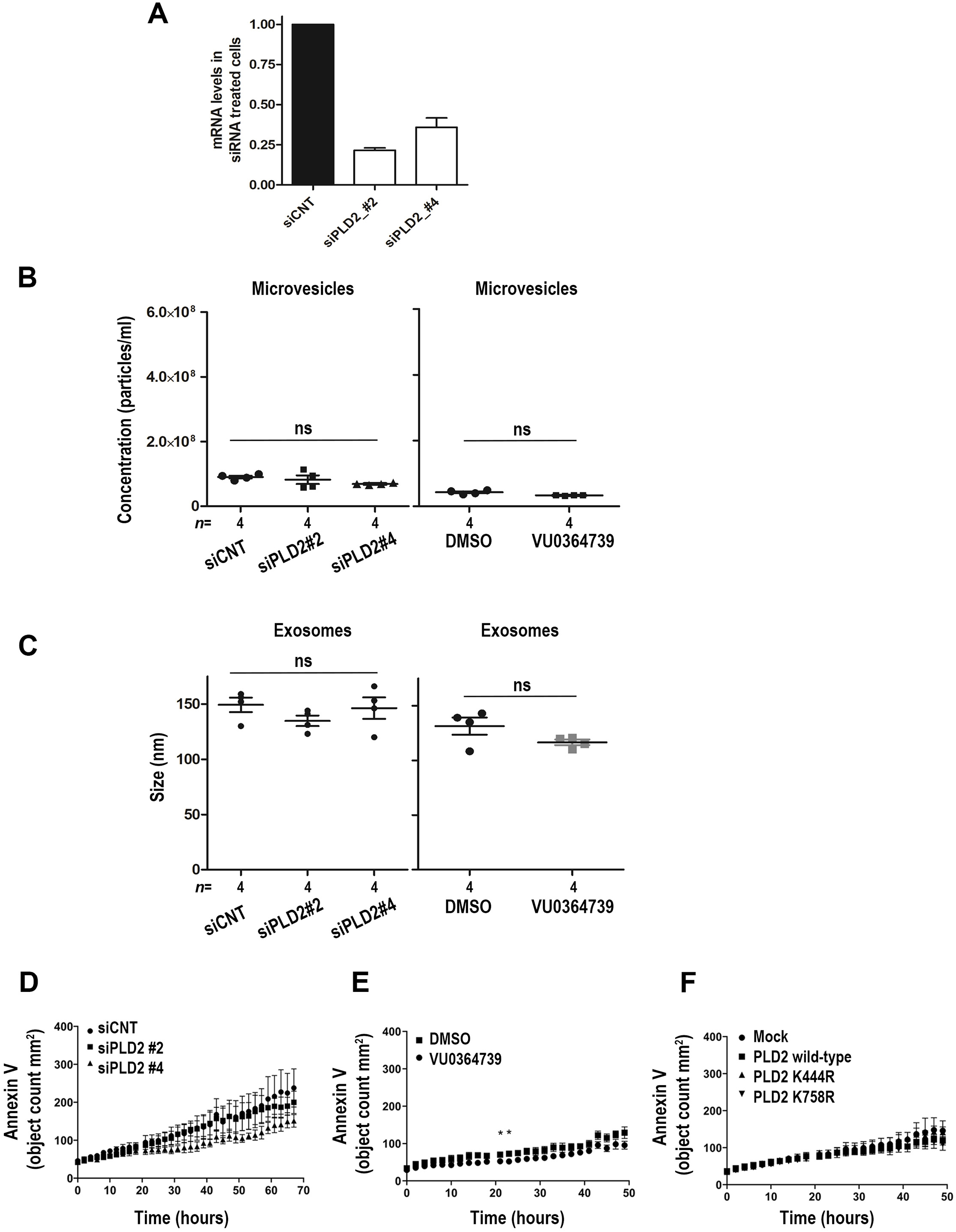
Related to Figure 1. Modulations of PLD2 lipase activity do not affect the number of microvesicles, the size of exosomes or the cell viability. **(A)** Effectiveness of the PLD2 RNAi treatments, using 2 different PLD2-specific RNAi sequences, as indicated. mRNA levels were expressed in arbitrary units and relative to two genes of reference (GAPDH and actin). **(B)** PLD2-loss effects on MVs, studied by NTA. MVs, isolated by ultracentrifugation (10K pellet), derived from control (siCNT) and PLD2-depleted MCF-7 cells, using two different PLD2-specific RNAi sequences (siPLD2#2 and siPLD2#4) (left panel), or derived from controls (DMSO) and cells treated with a specific PLD2 inhibitor (VU0364739) (right panel). The dot plot represents the total number of particles, with each independent experiment represented by a different symbol. NTA data shows that PLD2-depletion and PLD2-inactivation does not affect the number of MVs secreted by the cell. **(C)** Dot plot related to Fig. 1a, illustrating that PLD2-depletion (siPLD2#2 and siPLD2#4) or PLD2-inactivation (VU0364739) does not significantly affect the size of secreted particles sedimenting as 100K pellets, comparing treated cells with their respective controls (siCNT and DMSO). “ns” non-significant (Student’s t-test). **(D-F)** Kinetics of cell death as detected with Annexin V (green objects per mm^2^) using an IncuCyte ZOOM. **(D)** MCF-7 cells were transfected with a non-targeting siRNA (siCNT) or two independent siRNA sequences targeting PLDZ2 (siPLD2 #2 and siPLD2 #4). **(E)** Cells were treated with the PLD2 inhibitor VU0364739 or 0,1% DMSO. **(F)** Cells were transfected with Myc-tagged *wild-type* or lipase-dead (K444R or K758R) mutant PLD2, and the respective control cells transfected with empty vector (mock) vector. Each data point represents 3 technical replicates ± S.D. Two way ANOVA tests were performed using GraphPad Prism 5 software, *P<0.05.

**Figure S2.**
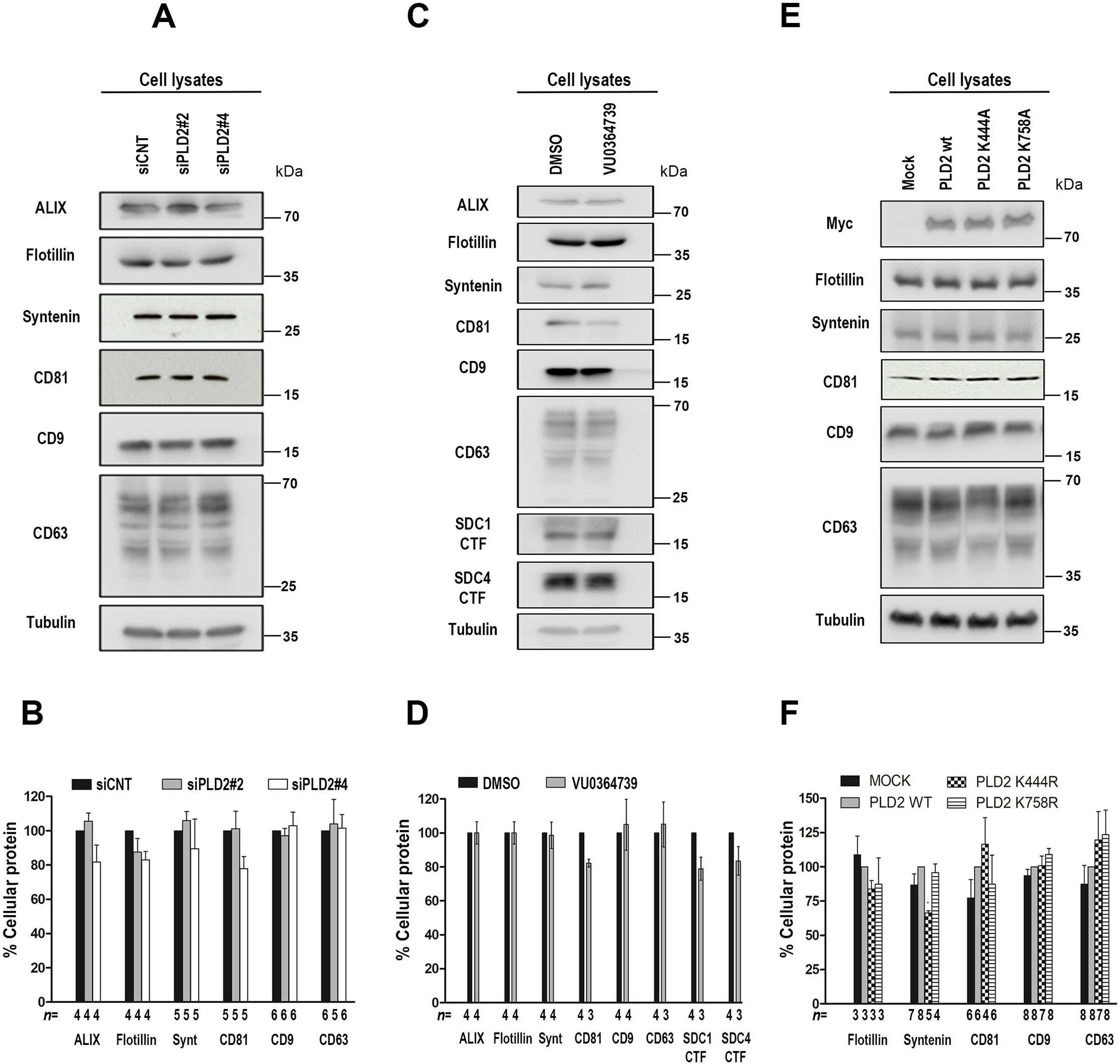
Related to Figure 2. Cellular levels of a selected set of markers proteins commonly enriched in exosomes are not affected after PLD2 loss-of-expression, inhibition of enzymatic activity, or gain-of-expression. **(A,B)** Western blot and histogram related to Fig. 2A,B illustrating that PLD2-depletion, in MCF-7 cells, does not significantly affect the cellular levels of the various marker proteins. **(C,D)** Western blot and histogram related to Fig. 2C,D illustrating that PLD2-inactivation, in MCF-7 cells, does not significantly affect the cellular levels of the various marker proteins. **(E,F)** Western blot and histogram related to Fig. 2E,F illustrating that gain of PLD2, whether *wild-type* or lipase-dead, in MCF-7 cells, does not significantly affect the cellular levels of the various marker proteins. Calculated for *n* independent experiments.

**Figure S3.**
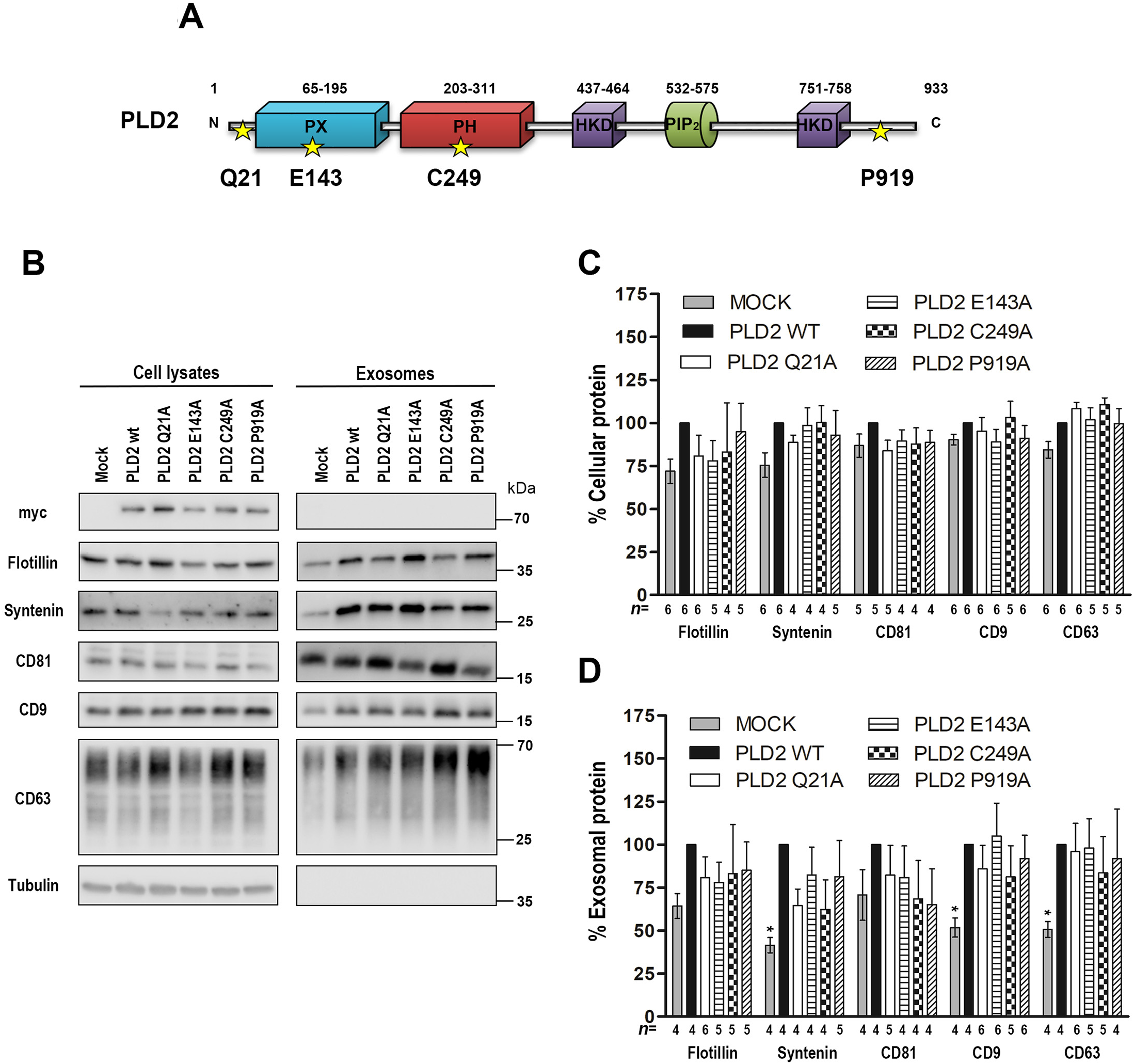
Related to Figure 2. PLD2 forms mutated in residues non-conserved during evolution do not affect the cellular or exosomal levels of a selected set of markers proteins commonly enriched in exosomes. **(A)** Domain structure of mammalian PLD2. The point mutations in residues that were not-conserved during evolution, used in this study as a control, are represented with yellow stars. UniProt code: PLD2 014393. **(B)** Exosomes secreted by MCF-7 cells overexpressing Myc-tagged *wild-type* or mutant PLD2 were characterized by western blot, using antibodies for several different markers, commonly found in EVs and enriched in exosomes in particular, as indicated (right panel). The gain of Myc-tagged PLD2, either *wild-type* or mutant, in MCF-7 cells, does not significantly affect the cellular levels of the various marker proteins (left panel). Histogram bars represent mean signal intensities ± SD in cell lysates **(C)** or exosomes **(D)**, relative to signals in lysates or exosomes from cells overexpressing *wild-type* PLD2, taken as 100 percent (black bars). Statistical tests were performed using GraphPad Prism 5 software. *P<0.05 (Student’s t-test), calculated for n independent experiments.

**Figure S4.**
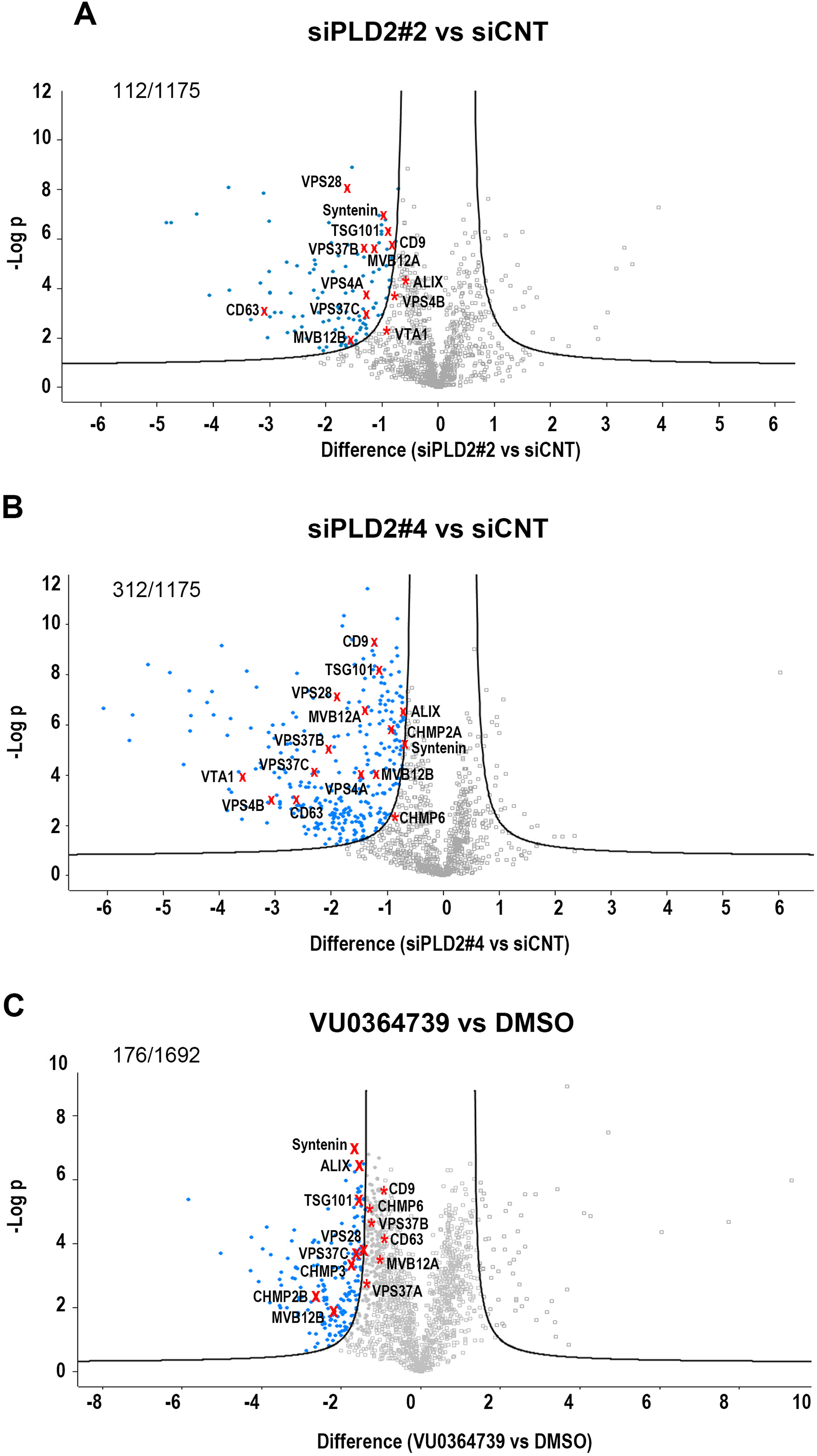
Related to Figure 4. Graph representation of ESCRT proteins identified in the proteomics analysis. Volcano Plots showing the significance two-sample t-test (-Log p value) vs. iBAQ intensity (Log2 (siPLD2#2 vs siCNT)) **(A)**, (Log2 (siPLD2#4 vs siCNT)) **(B)** or (Log2 (VU0364739 vs DMSO)) **(C)** on the y and x axes, respectively. Data result from three different experiments (n=2 for PLD2-depletion, n=1 for PLD2-inactivation) processed three times. ESCRT subunits, syntenin, ALIX, CD63 and CD9 are represented with red crosses or asterisks if they are affected in the proteomics analysis or not respectively. Proteins with significant or non-significant changes are represented with blue or grey dots, respectively. 1175 proteins were identified after PLD2-depletion and 112 **(A)** or 312 **(B)** were significantly decreased in exosomes secreted from MCF-7 cells treated with siPLD2#2 or siPLD2#4 respectively compared to control condition (siCNT). 1692 proteins were identified after PLD2-inactivation and 176 were significantly decreased in these exosomes compared to control condition (DMSO) **(C)**.

**Figure S5.**
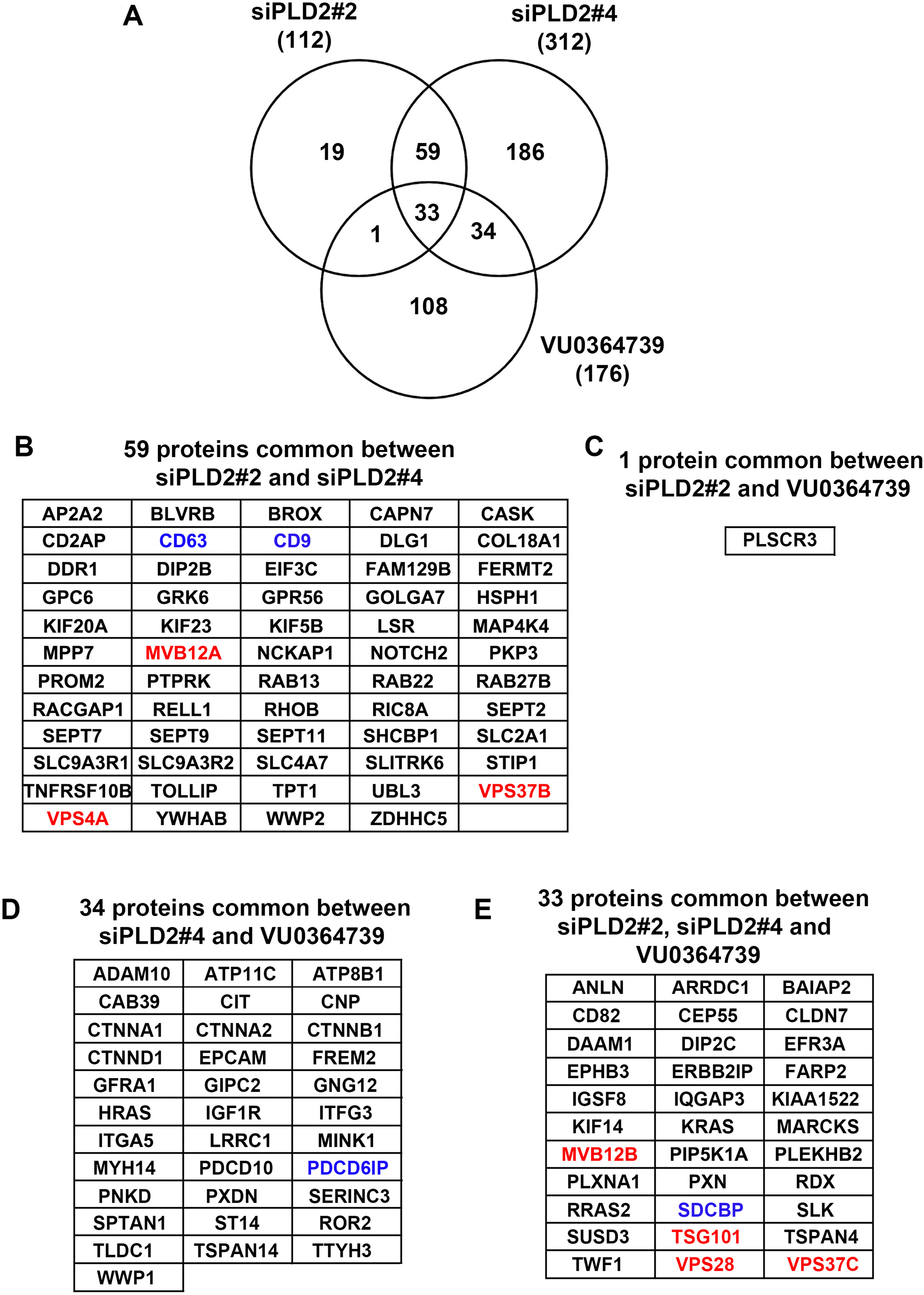
Related to Figure 4. List of proteins commonly affected in the proteomics analysis between different conditions. **(A)** Exosomes derived from control (siCNT) and PLD2-depleted MCF-7 cells, the latter treated with two different PLD2-specific RNAi sequences (siPLD2#2 and siPLD2#4), and exosomes derived from control cells (DMSO) and cells treated with a specific PLD2 inhibitor (VU0364739) were isolated for proteomics studies. A Venn diagram using overlapping circles is used to illustrate the relationship between the three different data set. Numbers in brackets correspond with the proteins significantly decreased in each condition, as indicated. Circles that overlap show the number of common proteins among groups. **(B**) List of 59 proteins commonly affected in the conditions siPLD2#2 vs siCNT and siPLD2#4 vs siCNT. **(C)** Protein commonly affected in the conditions siPLD2#2 vs siCNT and VU0364739 vs DMSO. **(D)** List of 34 proteins commonly affected in the conditions siPLD2#4 vs siCNT and VU0364739 vs DMSO. **(E)** List of 33 proteins commonly affected in the three conditions (siPLD2#2 vs siCNT, siPLD2#4 vs siCNT and VU0364739 vs DMSO). Proteins names of ESCRT members are labelled in red. Protein markers commonly found in exosomes and also used here for western blot analysis are labelled in blue.

**Figure S6.**
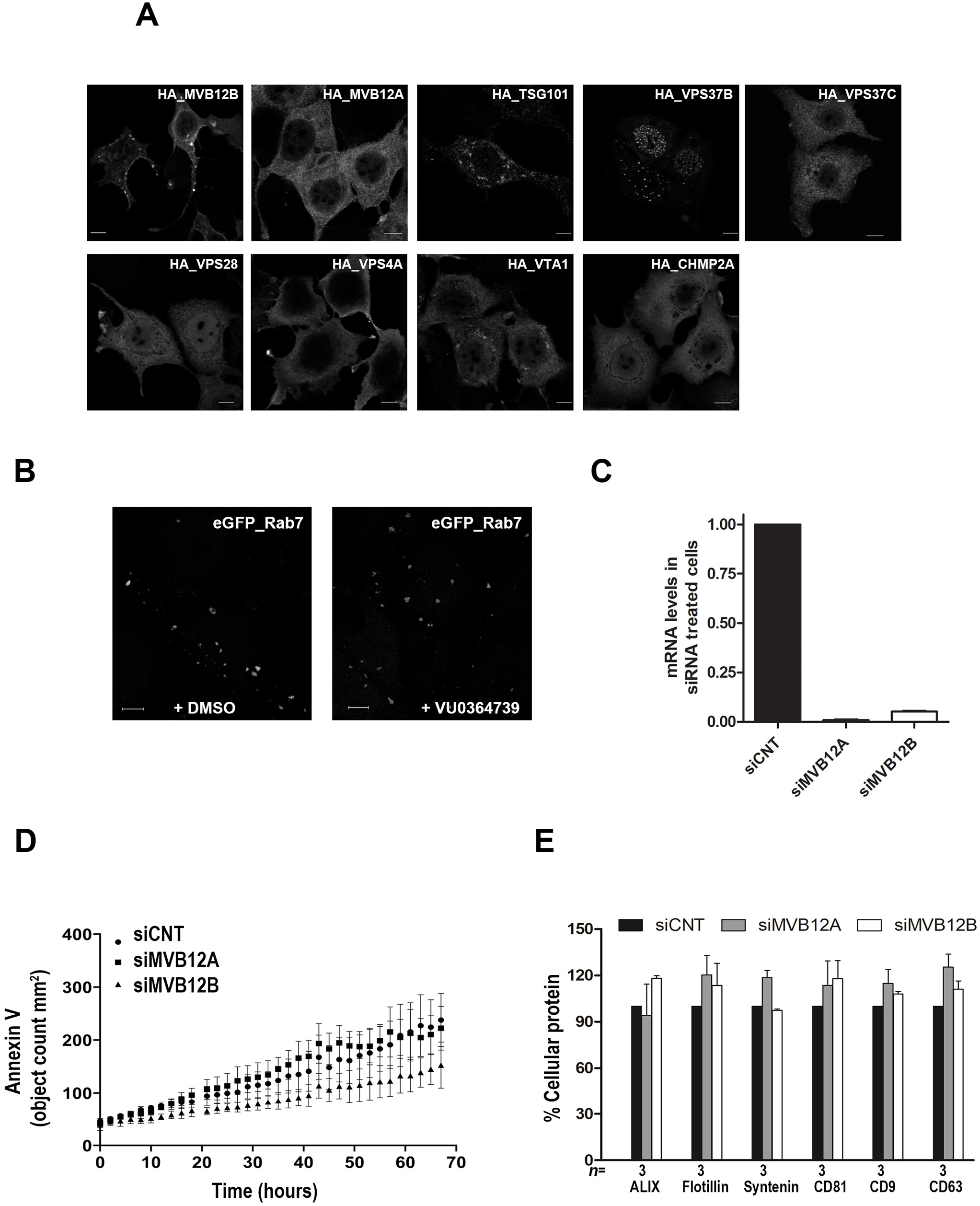
Related to Figure 5. Localization of HA-tagged ESCRT proteins, efficiency of MVB12 siRNAs and evidence that loss of MVB12A or MVB12B does not affect cell viability. **(A)** Representative confocal micrographs of MCF-7 cells showing the distribution of HA-tagged ESCRT proteins, as indicated. Note that HA-tagged (HA_) proteins have the same subcellular localization as the m-Cherry tagged proteins. Scale bar, 10 μm. **(B)** Effectiveness of the pools of four RNAi for MVB12A (siMVB12A) or MVB12B (siMVB12B), as indicated. mRNA levels were expressed in arbitrary units, and relative to two genes of reference (GAPDH and actin). **(C)** Kinetics of cell death of MCF-7 cells transfected with a pool of four RNAi for MVB12A (siMVB12A) or MVB12B (siMVB12B) or with non-targeting control RNAi (siCNT), as detected with Annexin V (green objects per mm^2^) using an IncuCyte ZOOM. Each data point represents 3 technical replicates ± S.D. Two-way ANOVA tests were performed using GraphPad Prism 5 software. **(D)** Histogram related to Fig. 5E, left panel, illustrating that MVB12A or MVB12B-depletion, in MCF-7 cells, does not significantly affect the cellular levels of the various marker proteins. Bars represent mean signal intensities ± SD in cell lysates, relative to signals measured in control sample (siCNT) (black bars), taken as 100%. Calculated for *n* independent experiments.

**Figure S7.**
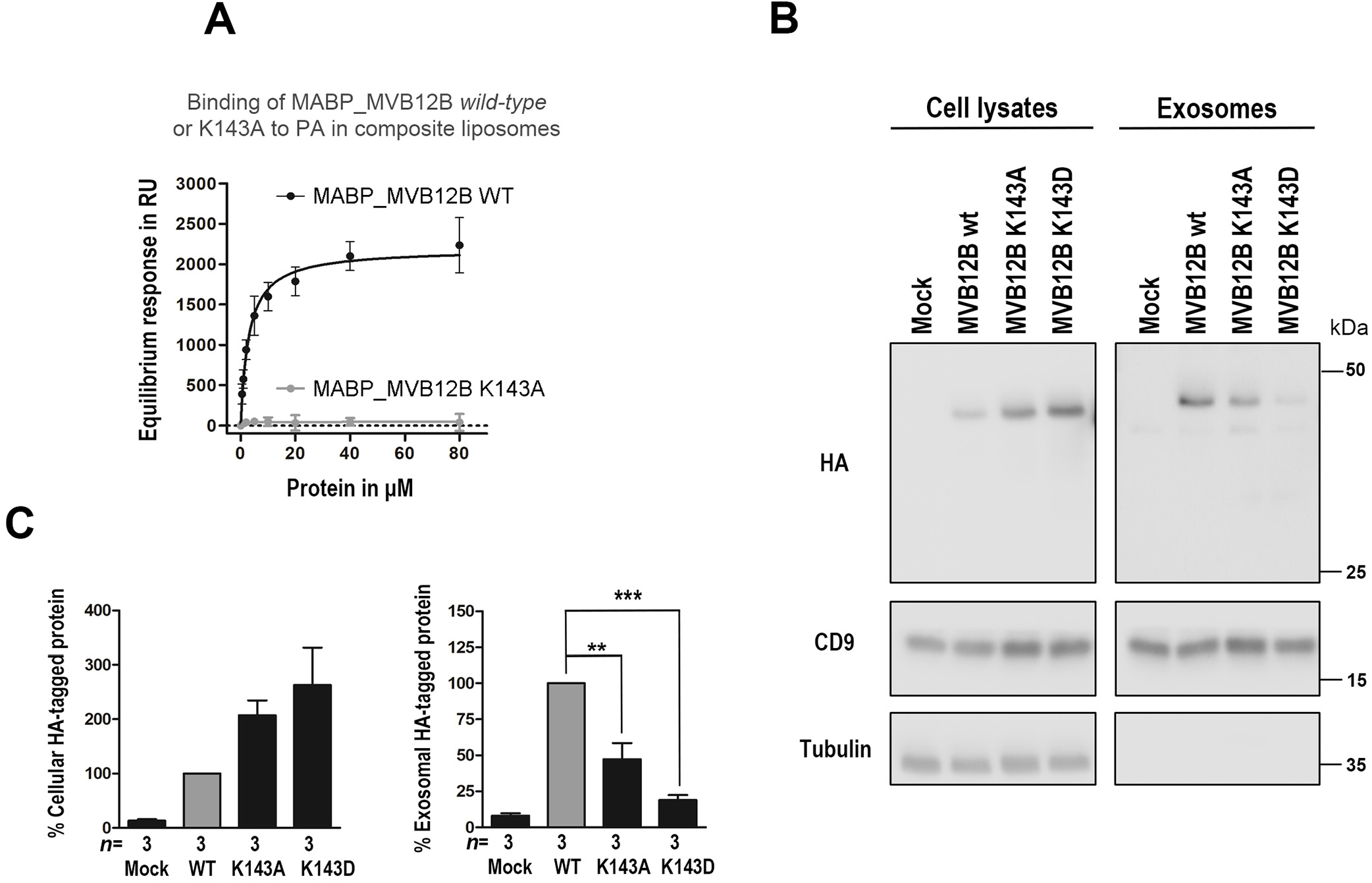
Related to Figure 6. MVB12B-PA binding mutants fail to be incorporated into exosomes. **(A)** Signals obtained at equilibrium using 2% PA-containing liposomes and *wild-type* MABP_MVB12B (black line) or MABP_MVB12B K143A (grey line), subtracted for association to 0%-PA liposomes and plotted as function of protein concentration. **(B)** Exosomes were purified by differential ultracentrifugation from the conditioned media of MCF-7 cells transiently transfected with HA-tagged MVB12B, either *wild-type* or mutant, or control cells (empty vector), as indicated. Total cell lysates (left panel) and the corresponding exosomes (right panel) were analyzed by western blotting using antibodies for several different markers, commonly found in EVs and enriched in exosomes in particular, as indicated. **(C)** Histograms representing mean signal intensities ± SD in cell lysates (left panel) or exosomes (right panel), relative to signals in lysates or exosomes from cells expressing wild-type MVB12, taken as 100% (grey bars). Statistical tests were performed using GraphPad Prism 5 software. **P<0.01, ***P<0.001 (Student’s t-test). Calculated for *n* independent experiments. Note that MVB12B *wild-type* is less abundant in cell lysates compared with the mutated forms K143A and K143D whereas the *wild-type* is more enriched in exosomes than the mutants.

## References

Babst, M. (2011). MVB vesicle formation: ESCRT-dependent, ESCRT-independent and everything in between. Current Opinion in Cell Biology 23, 452–457.

Bache, K.G., Brech, A., Mehlum, A., and Stenmark, H. (2003). Hrs regulates multivesicular body formation via ESCRT recruitment to endosomes. Journal of Cell Biology 162, 435–442.

Baietti, M.F., Zhang, Z., Mortier, E., Melchior, A., Degeest, G., Geeraerts, A., Ivarsson, Y., Depoortere, F., Coomans, C., Vermeiren, E., et al. (2012). Syndecan-syntenin-ALIX regulates the biogenesis of exosomes. Nature Cell Biology 14, 677–685.

Boura, E., and Hurley, J.H. (2012). Structural basis for membrane targeting by the MVB12-associated beta-prism domain of the human ESCRT-I MVB12 subunit. Proceedings of the National Academy of Sciences of the United States of America 109, 1901–1906.

Carlton, J.G., and Martin-Serrano, J. (2007). Parallels between cytokinesis and retroviral budding: A role for the ESCRT machinery. Science 316, 1908–1912.

Charrin, S., Le Naour, F., Oualid, M., Billard, M., Faure, G., Hanash, S.M., Boucheix, C., and Rubinstein, E. (2001). The major CD9 and CD81 molecular partner-Identification and characterization of the complexes. Journal of Biological Chemistry 276, 14329–14337.

Christ, L., Raiborg, C., Wenzel, E.M., Campsteijn, C., and Stenmark, H. (2017). Cellular Functions and Molecular Mechanisms of the ESCRT Membrane-Scission Machinery. Trends in Biochemical Sciences 42, 42–56.

Curtiss, M., Jones, C., and Babst, M. (2007). Efficient cargo sorting by ESCRT-I and the subsequent release of ESCRT-I from multivesicular bodies requires the subunit Mvb12. Molecular Biology of the Cell 18, 636–645.

de Souza, R.F., and Aravind, L. (2010). UMA and MABP domains throw light on receptor endocytosis and selection of endosomal cargoes. Bioinformatics 26, 1477–1480.

Egea-Jimenez, A.L., Gallardo, R., Garcia-Pino, A., Ivarsson, Y., Wawrzyniak, A.M., Kashyap, R., Loris, R., Schymkowitz, J., Rousseau, F., and Zimmermann, P. (2016). Frizzled 7 and PIP2 binding by syntenin PDZ2 domain supports Frizzled 7 trafficking and signalling. Nature Communications 7.

Egea-Jimenez, A.L., and Zimmermann, P. (2018). Phospholipase D and phosphatidic acid in the biogenesis and cargo loading of extracellular vesicles. Journal of Lipid Research 59, 1554–1560.

Gatta, A.T., and Carlton, J.G. (2019). The ESCRT-machinery: closing holes and expanding roles. Current Opinion in Cell Biology 59, 121–132.

Ghossoub, R., Chéry, M., Audebert, S., Leblanc, R., Egea-Jimenez, A.L., Lembo, F., Mammar, S., Le Dez, F., Camoin, L., Borg, J.-P., et al. (2020). Tetraspanin-6 negatively regulates exosome production. Proceedings of the National Academy of Sciences 117, 5913.

Ghossoub, R., Lembo, F., Rubio, A., Gaillard, C.B., Bouchet, J., Vitale, N., Slavik, J., Machala, M., and Zimmermann, P. (2014). Syntenin-ALIX exosome biogenesis and budding into multivesicular bodies are controlled by ARF6 and PLD2. Nature Communications 5.

Hanson, P.I., and Cashikar, A. (2012). Multivesicular Body Morphogenesis. Annual Review of Cell and Developmental Biology, Vol 28 28, 337–362.

Henne, W.M., Buchkovich, N.J., and Emr, S.D. (2011). The ESCRT Pathway. Developmental Cell 21, 77–91.

Kalluri, R., and LeBleu, V.S. (2020). The biology, function, and biomedical applications of exosomes. Science 367, 640–+.

Katzmann, D.J., Stefan, C.J., Babst, M., and Emr, S.D. (2003). Vps27 recruits ESCRT machinery to endosomes during MVB sorting. Journal of Cell Biology 162, 413–423.

Kooijman, E.E., Chupin, V., de Kruijff, B., and Burger, K.N.J. (2003). Modulation of membrane curvature by phosphatidic acid and lysophosphatidic acid. Traffic 4, 162–174.

Kostelansky, M.S., Schluter, C., Tam, Y.Y.C., Lee, S., Ghirlando, R., Beach, B., Conibear, E., and Hurley, J.H. (2007). Molecular architecture and functional model of the complete yeast ESCRT-I heterotetramer. Cell 129, 485–498.

Laulagnier, K., Grand, D., Dujardin, A., Hamdi, S., Vincent-Schneider, H., Lankar, D., Salles, J.P., Bonnerot, C., Perret, B., and Record, M. (2004). PLD2 is enriched on exosomes and its activity is correlated to the release of exosomes. Febs Letters 572, 11–14.

Lavieri, R.R., Scott, S.A., Selvy, P.E., Kim, K., Jadhav, S., Morrison, R.I., Daniels, J.S., Brown, H.A., and Lindsley, C.W. (2010). Design, Synthesis, and Biological Evaluation of Halogenated N-(2-(4-Oxo-1-phenyl-1,3,8-triazaspiro 4.5 decan-8-yl)ethyl)benzamides: Discovery of an Isoform-Selective Small Molecule Phospholipase D2 Inhibitor. Journal of Medicinal Chemistry 53, 6706–6719.

Lotvall, J., Hill, A.F., Hochberg, F., Buzas, E.I., Di Vizio, D., Gardiner, C., Gho, Y.S., Kurochkin, I.V., Mathivanan, S., Quesenberry, P., et al. (2014). Minimal experimental requirements for definition of extracellular vesicles and their functions: a position statement from the International Society for Extracellular Vesicles. Journal of extracellular vesicles 3, 26913–26913.

Mahankali, M., Henkels, K.M., Alter, G., and Gomez-Cambronero, J. (2012). Identification of the Catalytic Site of Phospholipase D2 (PLD2) Newly Described Guanine Nucleotide Exchange Factor Activity. Journal of Biological Chemistry 287, 41417–41431.

Margolis, L., and Sadovsky, Y. (2019). The biology of extracellular vesicles: The known unknowns. PLoS Biol 17, e3000363.

Marsh, M., and van Meer, G. (2008). Cell biology-No ESCRTs for exosomes. Science 319, 1191–1192.

Mathieu, M., Martin-Jaular, L., Lavieu, G., and Thery, C. (2019). Specificities of secretion and uptake of exosomes and other extracellular vesicles for cell-to-cell communication. Nat Cell Biol 21, 9–17.

Matsuo, H., Chevallier, J., Mayran, N., Le Blanc, I., Ferguson, C., Faure, J., Blanc, N.S., Matile, S., Dubochet, J., Sadoul, R., et al. (2004). Role of LBPA and Alix in multivesicular liposome formation and endosome organization. Science 303, 531–534.

Morita, E., Sandrin, V., Alam, S.L., Eckert, D.M., Gygi, S.P., and Sundquist, W.I. (2007). Identification of human MVB12 proteins as ESCRT-I Subunits that function in HIV budding. Cell Host & Microbe 2, 41–53.

Muralidharan-Chari, V., Clancy, J., Plou, C., Romao, M., Chavrier, P., Raposo, G., and D’Souza-Schorey, C. (2009). ARF6-Regulated Shedding of Tumor Cell-Derived Plasma Membrane Microvesicles. Current Biology 19, 1875–1885.

Oestreich, A.J., Davies, B.A., Payne, J.A., and Katzmann, D.J. (2007). Mvb12 is a novel member of ESCRT-I involved in cargo selection by the multivesicular body pathway. Molecular Biology of the Cell 18, 646–657.

Pang, S.S., Bayly-Jones, C., Radjainia, M., Spicer, B.A., Law, R.H.P., Hodel, A.W., Parsons, E.S., Ekkel, S.M., Conroy, P.J., Ramm, G., et al. (2019). The cryo-EM structure of the acid activatable pore-forming immune effector Macrophage-expressed gene 1. Nature Communications 10.

Pashkova, N., and Piper, R.C. (2012). UBAP1: A New ESCRT Member Joins the cl_Ub. Structure 20, 383–385.

Schoneberg, J., Lee, I.H., Iwasa, J.H., and Hurley, J.H. (2017). Reverse-topology membrane scission by the ESCRT proteins. Nature Reviews Molecular Cell Biology 18, 5–17.

Skotland, T., Ekroos, K., Kauhanen, D., Simolin, H., Seierstad, T., Berge, V., Sandvig, K., and Llorente, A. (2017). Molecular lipid species in urinary exosomes as potential prostate cancer biomarkers. European Journal of Cancer 70, 122–132.

Skotland, T., Hessvik, N.P., Sandvig, K., and Llorente, A. (2019). Thematic Review Series: Exosomes and Microvesicles: Lipids as Key Components of their Biogenesis and Functions Exosomal lipid composition and the role of ether lipids and phosphoinositides in exosome biology. Journal of Lipid Research 60, 9–18.

Slagsvold, T., Aasland, R., Hirano, S., Bache, K.G., Raiborg, C., Trambaiolo, D., Wakatsuki, S., and Stenmark, H. (2005). Eap45 in mammalian ESCRT-II binds ubiquitin via a phosphoinositide-interacting GLUE domain. Journal of Biological Chemistry 280, 19600–19606.

Stefani, F., Zhang, L., Taylor, S., Donovan, J., Rollinson, S., Doyotte, A., Brownhill, K., Bennion, J., Pickering-Brown, S., and Woodman, P. (2011). UBAP1 Is a Component of an Endosome-Specific ESCRT-I Complex that Is Essential for MVB Sorting. Current Biology 21, 1245–1250.

Strack, B., Calistri, A., Craig, S., Popova, E., and Gottlinger, H.G. (2003). AIP1/ALIX is a binding partner for HIV-1 p6 and EIAV p9 functioning in virus budding. Cell 114, 689–699.

Teo, H.L., Gill, D.J., Sun, J., Perisic, O., Veprintsev, D.B., Vallis, Y., Emr, S.D., and Williams, R.L. (2006). ESCRT-I core and ESCRT-II GLUE domain structures reveal role for GLUE in linking to ESCRT-I and membranes. Cell 125, 99–111.

Thery, C., Witwer, K.W., Aikawa, E., Alcaraz, M.J., Anderson, J.D., Andriantsitohaina, R., Antoniou, A., Arab, T., Archer, F., Atkin-Smith, G.K., et al. (2018). Minimal information for studies of extracellular vesicles 2018 (MISEV2018): a position statement of the International Society for Extracellular Vesicles and update of the MISEV2014 guidelines. Journal of Extracellular Vesicles 7.

Trajkovic, K., Hsu, C., Chiantia, S., Rajendran, L., Wenzel, D., Wieland, F., Schwille, P., Bruegger, B., and Simons, M. (2008). Ceramide triggers budding of exosome vesicles into multivesicular Endosomes. Science 319, 1244–1247.

van Niel, G., D’Angelo, G., and Raposo, G. (2018). Shedding light on the cell biology of extracellular vesicles. Nat Rev Mol Cell Biol 19, 213–228.

Vizcaino, J.A., Deutsch, E.W., Wang, R., Csordas, A., Reisinger, F., Rios, D., Dianes, J.A., Sun, Z., Farrah, T., Bandeira, N., et al. (2014). ProteomeXchange provides globally coordinated proteomics data submission and dissemination. Nature Biotechnology 32, 223–226.

Voelker, D.R. (1991). ORGANELLE BIOGENESIS AND INTRACELLULAR LIPID TRANSPORT IN EUKARYOTES. Microbiological Reviews 55, 543–560.

Welti, R., Li, W.Q., Li, M.Y., Sang, Y.M., Biesiada, H., Zhou, H.E., Rajashekar, C.B., Williams, T.D., and Wang, X.M. (2002). Profiling membrane lipids in plant stress responses-Role of phospholipase D alpha in freezing-induced lipid changes in Arabidopsis. Journal of Biological Chemistry 277, 31994–32002.

Whitley, P., Reaves, B.J., Hashimoto, M., Riley, A.M., Potter, B.V.L., and Holman, G.D. (2003). Identification of mammalian Vps24p as an effector of phosphatidylinositol 3,5-bisphosphate-dependent endosome compartmentalization. Journal of Biological Chemistry 278, 38786–38795.

Wunderley, L., Brownhill, K., Stefani, F., Tabernero, L., and Woodman, P. (2014). The molecular basis for selective assembly of the UBAP1-containing endosome-specific ESCRT-I complex. Journal of Cell Science 127, 663–672.

Yanez-Mo, M., Siljander, P.R.M., Andreu, Z., Zavec, A.B., Borras, F.E., Buzas, E.I., Buzas, K., Casal, E., Cappello, F., Carvalho, J., et al. (2015). Biological properties of extracellular vesicles and their physiological functions. Journal of Extracellular Vesicles 4.

